# Harnessing the CD2 axis to broaden and enhance the efficacy of CAR T cell therapies

**DOI:** 10.1101/2025.09.24.677515

**Authors:** Alberto Carturan, Mathew G. Angelos, Puneeth Guruprasad, Ruchi P. Patel, Raymone Pajarillo, Andrew Lee, Yunlin Zhang, Yi-Hao Chiang, Wei Xie, Jesse L. Rodriguez, Jaryse Harris, Pooja Devi, Olabisi I. Afolayan-Oloye, Jason Xu, Jonathan H. Sussman, Omar Elghawy, Austin Yang, Adam Barsouk, Jong Hyun Cho, Carolyn E. Shaw, Ekta Singh, Ositadimma Ugwuanyi, David Espie, Luca Paruzzo, Federico Stella, Shan Liu, Siena Nason, Antonio Imparato, Antonia Rotolo, Jean Lemoine, David M. Barrett, Avery Posey, Alain H. Rook, Vinodh Pillai, Adam Bagg, Stefano A. Pileri, Dongfang Liu, Kai Tan, Stephen J. Schuster, David T. Teachey, Patrizia Porazzi, Marco Ruella

## Abstract

Patients with T-cell lymphomas and leukemias have overall poor outcomes due to the lack of targeted and effective treatments, particularly in the relapsed and refractory settings. Development of chimeric antigen receptor (CAR) T-cells against T-cell neoplasms is limited by a lack of discriminating T-cell antigens that allow for effective anti-tumor responses while preventing CAR T-cell fratricide. We hypothesized that targeting CD2, a pan-T-cell antigen, using anti-CD2 CAR T-cells engineered without CD2 expression (CART2), would support CAR T-cell manufacturability and preclinical efficacy. Optimized CD2-knockout CART2, generated using CRISPR-Cas9, eradicated primary patient-derived CD2+ hematological neoplasms in vitro and in vivo, secreted effector cytokines, and exhibited adequate proliferative capacity. Nevertheless, CD2 has a key costimulatory function, and its deletion could lead to CAR T-cell dysfunction. Therefore, we tested the role of the CD2:CD58 axis in CAR T-cells, using the anti-CD19 CART models. We demonstrate that CD2 loss attenuates CART19 efficacy by reducing avidity for tumor antigen, co-stimulation, and ultimately in vivo activity. Analogously, we show that tumor CD58 loss reduces CART19 efficacy. To overcome this issue, we developed a novel PD-1:CD2 switch receptor that rescues intracellular CD2 signaling, particularly when PD-L1 is engaged, resulting in improved in vivo outcomes. Collectively, we studied the role of CD2 both as a target for CAR T cell therapy and as a critical costimulatory protein, whose signaling can be rescued using the PD-1:CD2 switch receptor. This receptor can be incorporated into CAR T-cells and provides an effective strategy to overcome CD2-signaling deficiencies.

## INTRODUCTION

Over the past decade, Chimeric antigen receptor (CAR) T-cell therapy has revolutionized the management of relapsed or refractory (r/r) B-cell malignancies(1–6). However, their application in T-cell leukemias and lymphomas remains restricted by fundamental immunobiological barriers, including the absence of a uniform target antigen and CAR T-cell fratricide leading to T-cell aplasia(7,8). Although several targets have been investigated, clinical progress has been hindered by heterogeneous antigen expression and insufficient *in vivo* expansion(9–13). Furthermore, this heterogeneity increases relapse risk by allowing persistence of malignant clones(8).

Among the most promising pan–T-cell targets are CD5 and CD7. To prevent fratricide(14,15), genetic and molecular strategies abrogating endogenous antigen expression on CAR T-cells have shown efficacy, with CD7-directed approaches representing the most advanced in clinical development(16–20). Early-phase clinical trials have demonstrated anti-CD7 CAR T-cells to be safe and preliminarily effective, with complete response rates of 80–100%(21–26). However, long-term follow-up indicates relapse in ∼30% of patients, partly due to CD7-antigen-negative escape(27), highlighting the need to investigate additional targets as alternatives, rescue therapies, or in combinatorial CAR T-cell strategies.

CD2 is a transmembrane glycoprotein of the immunoglobulin supergene family expressed on mature T-, NK-, and T-cell progenitors(28), as well as across T-cell malignancies, positioning it as an attractive target for cellular immunotherapies. CD2 interacts with CD58 (LFA-3) on antigen-presenting and target cells(29–31), in concert with molecules such as PD-1:PD-L1, to facilitate immune synapse formation between TCR and MHC(32–34). Intracellularly, CD2 provides costimulation that promotes T-cell activation, proliferation, and effector function in an MHC-dependent manner(35–37). Loss of CD2:CD58 interactions is associated with immune evasion and poor anti-tumor responses(38). However, since CAR signaling is MHC-independent, the role of CD2 in CAR T-cell immunobiology remains undefined.

We hypothesized that anti-CD2 CAR T-cells with endogenous CD2 knock-out (CART2) would be highly efficacious against CD2-expressing neoplasms. We selected a lead anti-CD2 CAR construct and, through a CRISPR-Cas9–based knock-out (KO) manufacturing process eliminating fratricide, generated a product with potent *in vitro* and *in vivo* efficacy against patient-derived T-cell leukemia. However, when using anti-CD19 CAR T-cells (CART19) against B-cell lymphoma, we observed that CD2 deletion in T-cells or CD58 deletion in tumor cells attenuated CAR T-cell activity. This deficiency was reversible with intracellular CD2 signaling rescue provided by a PD-1:CD2 switch receptor engineered into CART19. Collectively, our data support clinical investigation of CART2 in patients with r/r T-cell neoplasms and delineate a critical role for the CD2-CD58 axis in CAR T-cell immunobiology.

## METHODS

### Full methods are provided in the Supplemental Materials

Nalm6, Jurkat, OCI-Ly18, HEK293T (ATCC), and the TH20 T-ALL PDX (39) were maintained in RPMI-based R10; all lines were STR-authenticated and mycoplasma-free. Primary Sézary cells were from Dr. Alain Rook, and AML/T-ALL samples were collected at Penn under IRB approval (#855418).Genome-edited CAR T cells were generated from CD4^+^/CD8^+^ T cells (1:1) from the Penn Human Immunology Core (IRB #705906), electroporated with Cas9–sgRNA RNPs targeting CD2 or CD58 (**Tables S1–S2**), cultured in IL-7/IL-15, activated with CD3/CD28 Dynabeads, and transduced with lentiviral CAR vectors (MOI 1). Murine T cells were engineered similarly, and lentiviruses were produced in HEK293T cells, concentrated, and designed with 4-1BB/CD3ζ signaling; the MEDI507 orientation was used for definitive studies (**Table S3**). CAR expression was detected by flow cytometry using anti-idiotype antibodies (CAR19) or anti-G4S linker antibodies (CAR2, murine CAR19); a list of specific antibodies is provided in the Supplemental Materials. Cytotoxicity was quantified by luciferase-based assays or CellTrace Violet dilution, while proliferation and metabolism were assessed by flow cytometry and Seahorse XF96 profiling. Avidity was measured with Z-Movi acoustofluidic chips (LUMICKS™). Jurkat triple-reporter assays (E:T 1:1) evaluated NFAT, NF-κB, and AP-1 activity after co-culture with CAR constructs ± full-length or truncated switch receptors. Imaging assays used supported lipid bilayers functionalized with biotinylated CD19 to stimulate CAR T cells, with synapse formation visualized on an Olympus FV3000 confocal microscope(40)or analyzed in NALM6-GFP co-cultures by Leica Stellaris. In vivo experiments were performed in NSG or BALB/c mice bearing human or murine tumors, with CAR T infusion after engraftment or cyclophosphamide conditioning, and disease monitored by IVIS or caliper measurements(41).Single-cell RNA sequencing was performed using 10X v2 chemistry, sequenced on NovaSeq, and analyzed in Seurat v5 with clusterProfiler(42–44). Clinical cohorts included pediatric/young adult T-ALL (CHOP, 2002– 2014), PTCL from the European Institute of Oncology, and CITE-seq biobanking studies(45–48) CD58 immunohistochemistry was scored by blinded hematopathologists.

## RESULTS

### CD2 is highly expressed in T-cell lymphomas and leukemias

To establish the robustness of CD2 as a cancer antigen, we retrospectively analyzed its surface expression using clinical samples of T-cell lymphomas and leukemias. High (>75%) CD2 expression was observed in 77/90 (85%) adult peripheral T-cell lymphoma (PTCL) patients by immunohistochemistry (**Fig. 1A**) and in 48/51 (94%) pediatric T-cell acute lymphoblastic leukemia (T-ALL) patients by flow cytometry (**Fig. 1B**). In PTCL, while most samples showed high CD2, only 12/92 (13%) biopsies had high CD7. Of note, in CD2+/CD7− PTCL cases, CD7 expression was restricted to reactive T-cells (**Fig. 1C**).

**FIGURE 1.**
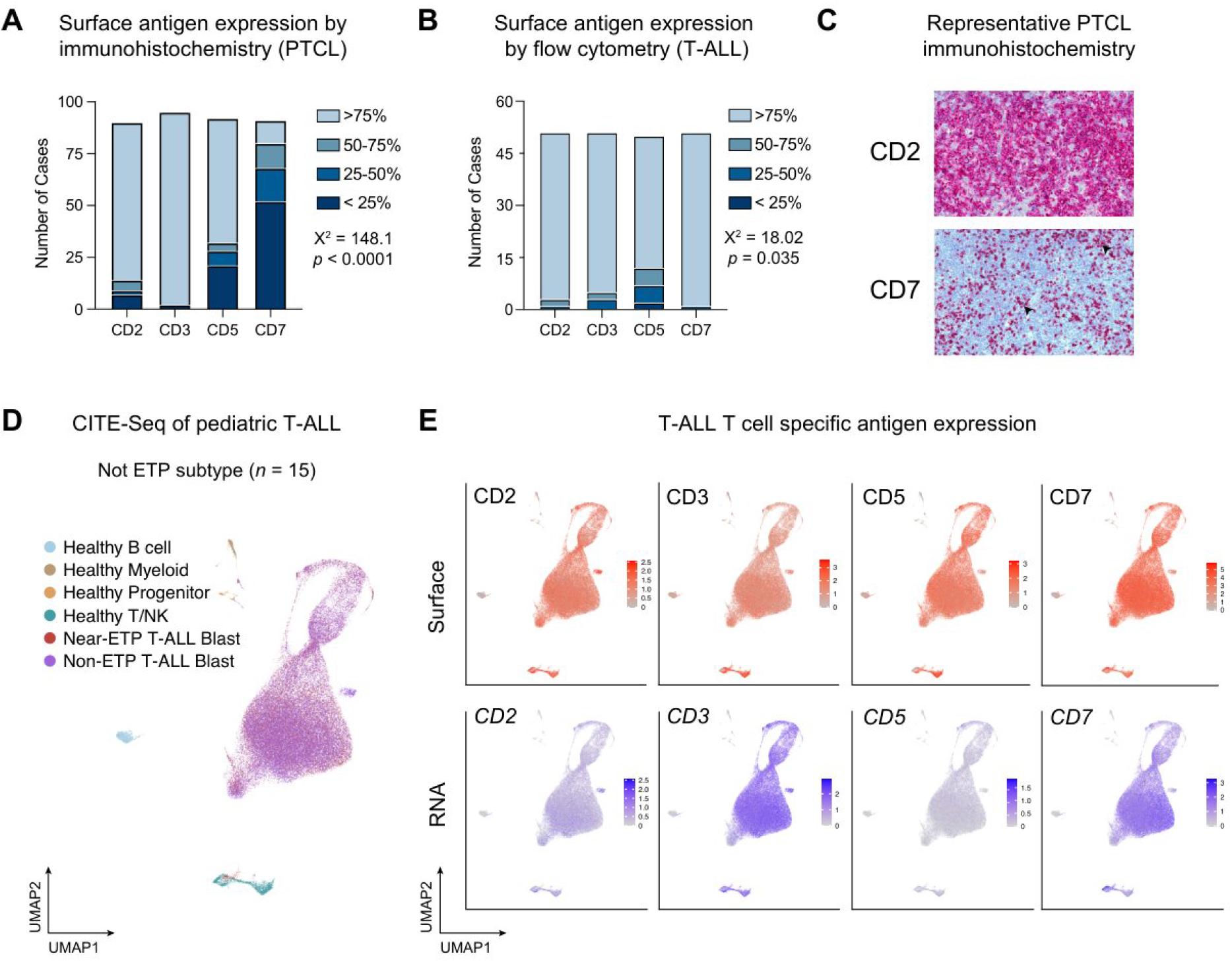
CD2 is highly expressed across pediatric and adult T-cell neoplasms. **(A)** Number of unique patients with CD2 expression in real-world, adult peripheral T-cell lymphoma (PTCL; n = 90), as determined by clinically validated immunohistochemistry. Percentage cohorts represent quantification of CD2 expression of the total biopsy. **(B)** Number of unique patients with surface CD2 expression in real-word, pediatric T-ALL (n = 51), as determined by clinically validated multiparameter flow cytometry and verified by a hematopathologist. Percentage cohorts represent quantification of total CD2 expression. Panels (A) and (B) were analyzed by categorical Chi-squared test. **(C)** Representative immunohistochemistry staining of a CD2^+^/CD7^-^ peripheral T-cell lymphoma. Arrowhead = reactive T-cells. **(D)** Diagnostic samples from 15 not-early T-cell precursor (ETP) T-cell acute lymphoblastic leukemia (T-ALL) patients (5 Near-ETP and 10 Non-ETP), treated on the phase 3 clinical trial AALL0434, were analyzed by CITE-seq as previously described by Tan K., *et al* (46). Shown is the UMAP representation of the complete dataset integrating transcriptomic and surface protein expression. **(E)** Analysis of pan–T-cell antigen expression from the CITE-seq dataset shown in (D). *Top*: surface protein expression; *Bottom*: RNA expression. Heat maps display normalized expression values; shading is limited to the 99th percentile for visualization purposes.

We next analyzed 40 pediatric T-ALL samples collected at diagnosis from the Children’s Oncology Group AALL0434 trial(47) using Cellular Indexing of Transcriptomes and Epitopes by sequencing (CITE-seq). This cohort, enriched with relapsed/refractory (r/r) cases as previously described(46), revealed CD2 surface and RNA expression in T-ALL blasts. Robust surface CD2 expression (>50% blasts) was detected in 73% of non-ETP T-ALL (11/15, n=8 r/r) (**Fig. 1D, 1E**) and in 40% of ETP-ALL (10/25, n=5 r/r) (**Fig. S1A**). The expression patterns noted in T-ALL patients closely matched their developmental arrest state; within developing thymocytes, we observed low, but detectable surface CD2 at the pro-T (CD34+CD1a-) stage, rising expression in pre-T (CD34+CD1a+) and double positive cell states (CD4+CD8+), and peak expression in functional alpha-beta T-cells (CD4/8 single positive, PTCRA-, TRAV/D/J+) (**Fig. S1B**). These data align with recent whole-genome subtyping of the AALL0434 cohort, where 74% (987/1335) of cases arrest in CD2^+^ state(49). Notably, in both T-ALL and normal thymocytes, surface CD2 expression persisted despite low/absent RNA. Collectively, CD2 emerges as a robust and homogeneously expressed target across T-cell developmental stages in both immature and mature neoplasms.

### Selection of a lead anti-CD2 CAR and development of a CRISPR-Cas9 CD2 knock-out manufacturing platform

Since CD2 is broadly expressed in T-cell leukemias and lymphomas, we sought to develop anti-CD2 CAR T-cells as a therapy for T-cell neoplasms. Given that CD2 is also expressed in healthy T-cells, leading to CAR T-cell fratricide, we established a strategy to delete CD2 during CAR T-cell manufacturing. First, we screened CRISPR-Cas9 single-guide RNAs (gRNAs) targeting *CD2*, identified the most efficient (gRNA08) by measuring CD2 protein loss in Jurkat cells (Fig. S2A), and validated it in primary human T-cells(50) (**Fig. 2A, 2B**).

**FIGURE 2.**
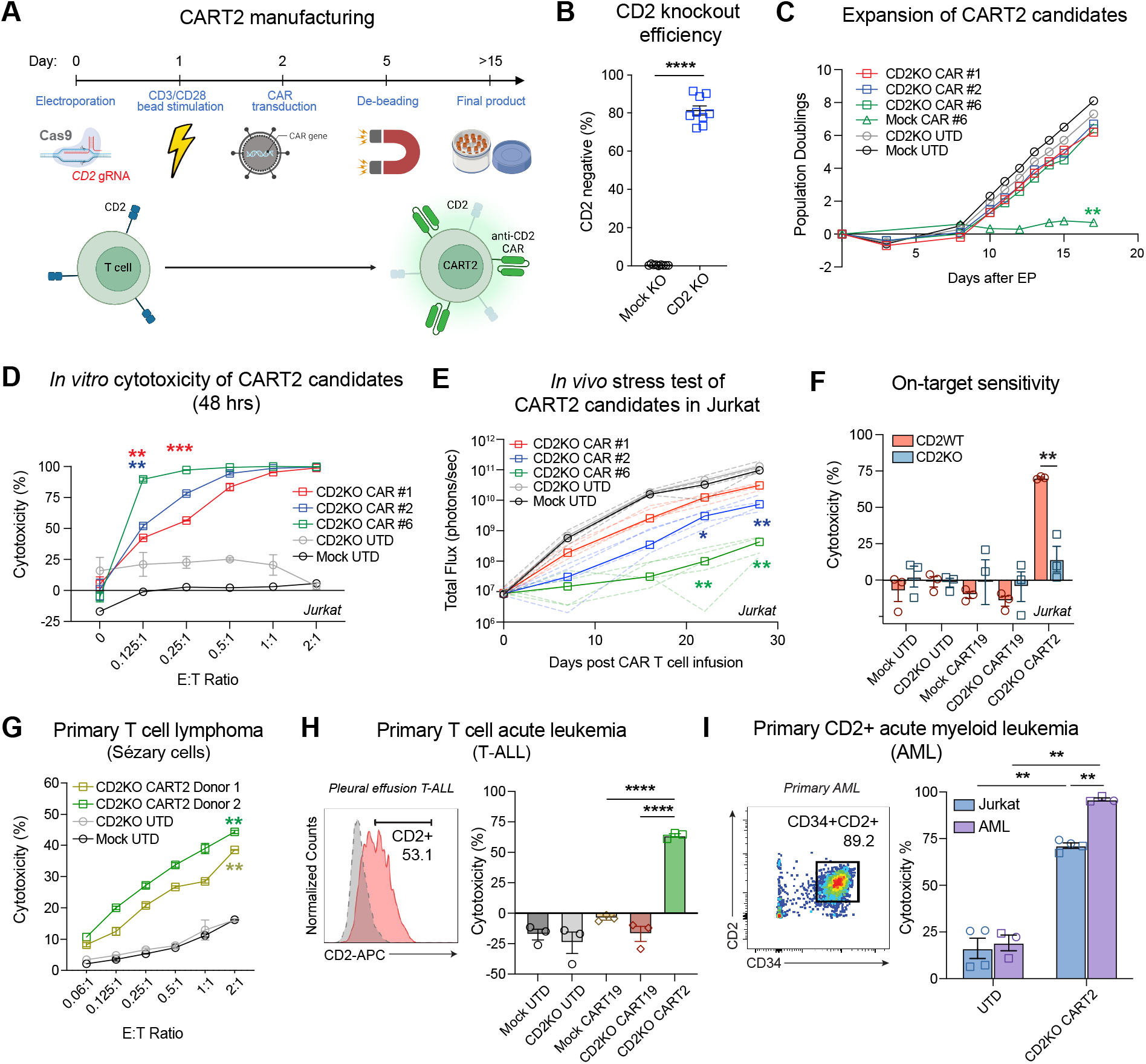
Anti-CD2 CAR T-cells (CART2) are manufacturable and specific against CD2-positive T-cell neoplasms. **(A)** CART2 expansion protocol schema. **(B)** Percentage of CD2-negative T-cells at Day 5 following CD2 gRNA electroporation and quantified by flow cytometry, n=9 independent healthy donors, analyzed by Student’s t-test. **(C)** Population doublings, as calculated by normalized viable cell count relative to total number of cells electroporated, of Mock UTD, CD2^KO^ UTD, and the top three CD2^KO^ CART2 candidates. Endogenous CD2 retention (Mock CAR #6) demonstrates a fratricide-limiting effect on CART2 expansion. **(D)** Percent cytotoxicity of the top three CD2^KO^ CART2 candidates against Jurkat cells after 48 hours of coculture. Significance is relative to CD2^KO^ CAR #6. **(E)** Bioluminescence imaging of tumor burden in each *NSG* mouse (n=4 mice per group) engrafted with a luciferase-positive CD2-positive T-ALL cell line (Jurkat) and treated with 1.0 x 10^6^ untransduced (UTD) or CD2^KO^ CAR-positive CART2 candidates (CAR #1, CAR #2, CAR #6). Bolded lines represent the median of each group. Representative data of 2 independent experiments are shown. Significance is relative to CD2^KO^ CAR #1. (**F)** Percent cytotoxicity of UTD, CART19, or CD2^KO^ CART2 cells against CD2-wild type (CD2^WT^) or CD2^KO^ Jurkat cells after 72 hours co-culture (E:T ratio = 0.5:1). **(G)** Percent cytotoxicity of Mock UTD, CD2^KO^ UTD, or CD2^KO^ CART2 cells against CD2-positive primary cutaneous T-cell lymphoma/Sezary cells after 24 hours coculture. Representative cytotoxicity curves from two independent CD2^KO^ CART2 products are shown. Significance is relative to CD2^KO^ UTD. CAR #6 **(H)** *Left*: Immunophenotype of a diagnostic malignant pleural effusion aspirate demonstrating low-level surface CD2 expression in T-cell blasts (red) as compared to non-malignant cells (grey). *Right*: Percent cytotoxicity of UTD, CART19, or CD2^KO^ CART2 cells against collected CD2-positive primary T-ALL after 72 hours of coculture (E:T ratio = 0.125:1). **(I)** CART2 effectively kills primary, patient-derived, CD2-expressing myeloblasts. *Left*: CD34 and CD2 expression of primary myeloblasts sorted from peripheral blood in a patient with a new diagnosis of acute myelogenous leukemia (AML). *Right*: Percent cytotoxicity of CD2^KO^ CART2 against control Jurkat cells and primary-patient derived CD2-positive AML from two independent donors (shown as either circle or square data points). Aside from Panel (B), all other panels were analyzed by ANOVA with Tukey post-hoc multiple comparisons test; **** p < 0.0001, *** p < 0.001, ** p < 0.01, * p < 0.05.

We next sought to generate anti-CD2 CAR constructs by screening six distinct single-chain variable fragments (scFv) derived from four CD2-antibodies (OKT11, T11.2, TS2_18.1.1, and MEDI507), tested in both heavy-to-light and light-to-heavy orientations (**Table S3**). Among them, three (CAR#1, CAR#2, and CAR#6) showed consistent expression >50% after cryopreservation (**Fig. S2B**) and were advanced for further development. CD2^KO^ CART2 expanded comparably to CD2 wild-type (CD2^WT^) untransduced T-cells (Mock UTD) and CD2^KO^ UTD T-cells, whereas CD2^WT^ anti-CD2 CAR T-cells, as expected, failed to expand due to fratricide (**Fig. 2C**).

We first evaluated the anti-tumor efficacy of the three lead CART2 constructs, all of which demonstrated potent cytotoxicity against CD2+ Jurkat cells *in vitro* (**Fig. 2D**). To identify the best-in-class construct, we used an *in vivo* stress model in which NOD-SCID-IL2Rg^null^ (*NSG*) mice were engrafted with Jurkat cells and randomized to receive Mock UTD, CD2^KO^ UTD, or one of the candidate CD2^KO^ CART2 products. CD2^KO^ CART2 engineered with a MEDI507-based scFv (CAR#6) most effectively controlled leukemia *in vivo* and was selected as the lead construct for all subsequent studies (**Fig. 2E**).

At the end of manufacturing, CD2^KO^ CART2 cells exhibited memory T-cell subset distributions comparable to UTD controls (CD2^WT^ and CD2^KO^) (**Fig. S2C**). We first confirmed that CD2^KO^ CART2 cells were able to secrete effector cytokines (i.e., IFN-γ, TNFα, IL-2) upon non-specific (PMA/Ionomycin) and antigen-specific (CD3/CD28 beads or CD2+ Jurkat) stimulation (**Fig. S2D**). CD2^KO^ CART2 proliferated in response to irradiated CD2+ Jurkat cells (**Fig. S2E**), unlike CD2^WT^ or CD2^KO^ UTD cells, while CD3/CD28 stimulation induced similar proliferation across groups (**Fig. S2F**).

### CD2-knockout CART2 is effective against CD2^+^ neoplasms in vitro and in vivo

We next evaluated the specificity and sensitivity of CD2^KO^ CART2 cells against CD2-expressing neoplasms *in vitro*. CD2^KO^ CART2 cells efficiently eliminated CD2^+^ Jurkat cells while showing no off-target activity against CD2^KO^ Jurkat (**Fig. 2F**) or CD2^−^ Nalm6 B-ALL controls (**Fig. S2G**). CD2^KO^ CART2 also demonstrated robust efficacy against primary T-cell lymphoma (Sézary cells) (**Fig. 2G**), primary T-ALL (**Fig. 2H**), and even patient-derived acute myeloid leukemia with aberrant CD2+ blast expression(51) (**Fig. 2I**). Collectively, these findings establish potent and specific anti-tumor activity of CD2-deleted CART2 across CD2-expressing neoplasms in both preclinical models and primary samples.

We then assessed *in vivo* efficacy of CD2^KO^ CART2 using patient-derived xenograft (*PDX*) T-ALL models. Primary lymphoblasts from a pediatric patient enrolled in the COG AALL1231 trial (NCT02112916)(39), were first transduced with luciferase for disease monitoring and engrafted into *NSG* mice. Then, mice were randomized to receive Mock UTD, CD2^KO^ UTD, or CD2^KO^ CART2 (**Fig. 3A, S3A**). CD2^KO^ CART2 effectively controlled systemic and central nervous system leukemia (**Fig. 3B, S3B**) and significantly prolonged survival compared with controls (**Fig. 3C**). These findings were reproduced using an independent donor **(Fig. S3C)**, further associated with robust peripheral blood expansion of CD2^KO^ CART2 **(Fig. S3D)** and a favorable safety profile **(Fig. S3E)**.

**FIGURE 3.**
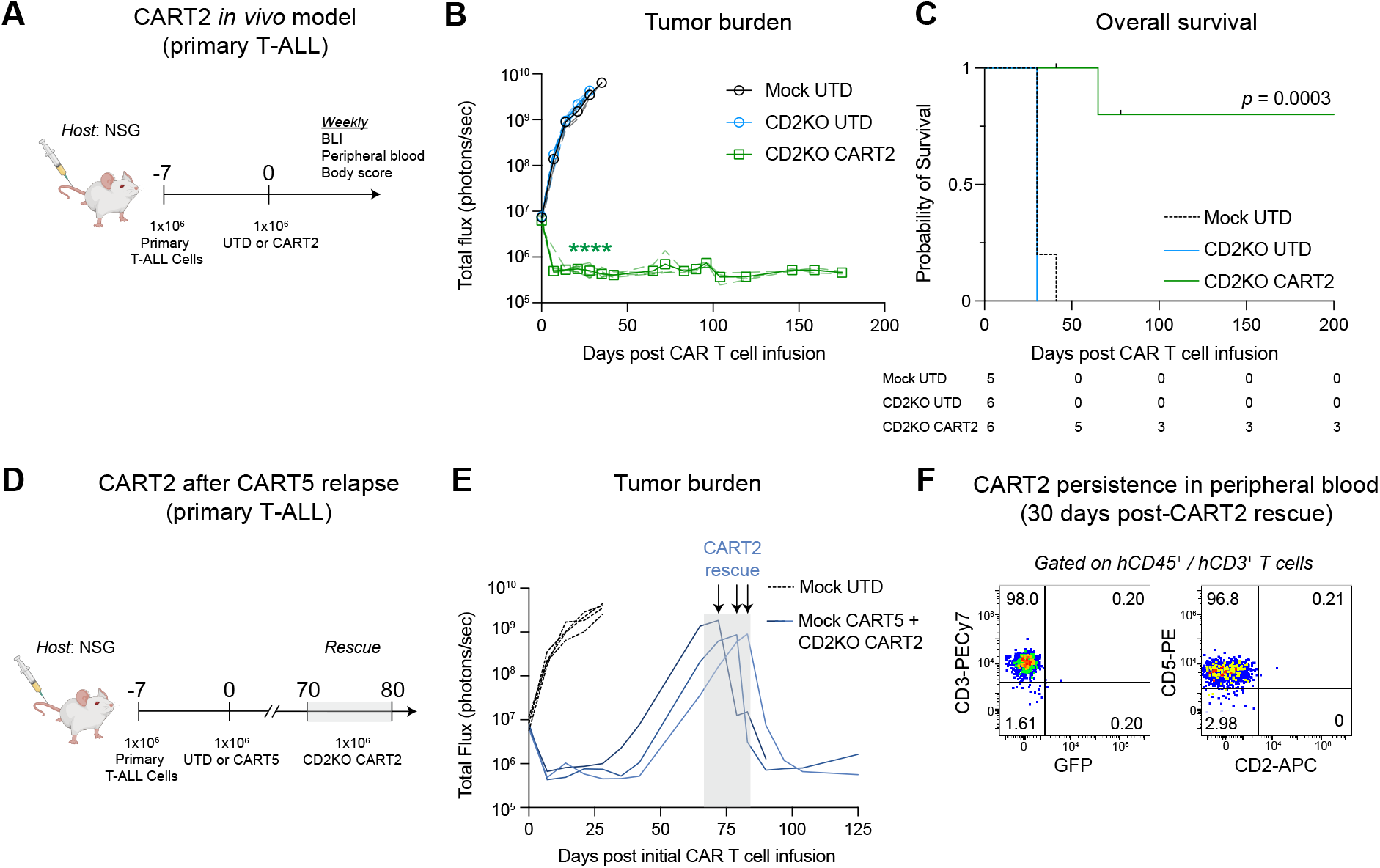
CART2 is highly effective *in vivo* against primary T-ALL. **(A)** Schema of CART2 treatment in patient-derived T-ALL xenografts (*PDX* T-ALL). *NSG* mice were engrafted with 1 x 10^6^ primary human T-ALL cells engineered to express GFP/Luciferase on day -7 and then injected with 1 x 10^6^ untransduced (Mock UTD), CD2^KO^ UTD, or CART2 on day 0. **(B)** Representative bioluminescence imaging of leukemic burden and **(C)** overall survival (median OS: CD2^KO^ CART2: not reached vs. Mock UTD: 30 days vs. CD2^KO^ UTD 30 days); n= 5-6 mice per group, additional CART2 donor is shown in Supplemental Figure 3. ANOVA with Tukey post-hoc multiple comparisons test was used for BLI analyses, **** p < 0.0001; Wilcox rank-sum test was performed for survival curve comparisons, p=0.0003. **(D)** Schema of CART2 rescue following CART5 relapse. Engrafted *PDX* T-ALL were injected with 1 x 10^6^ Mock UTD or Mock CART5 at day 0. 1 x 10^6^ CART2 cells were then injected in relapsed mice as retreatment upon disease relapse at high leukemia burden between Days 70-80. **(E)** Bioluminescence imaging of leukemic burden in each *PDX* T-ALL mouse (n=3 per group). **(F)** Representative flow cytometry plots of dual-positive human CD45+/human CD3+ T-cells from mouse peripheral blood collected at Day 30 following CART2 rescue. GFP = Green fluorescent protein.

We then developed a T-ALL model that was resistant to non-edited (CD5^WT^) anti-CD5 CAR T-cells (CART5), a target currently under clinical investigation for T-cell malignancies. Here, we treated *PDX* T-ALL mice with a suboptimal dose of non-edited CART5 to foster leukemic relapse within 100 days (**Fig. 3D**). Upon high leukemic burden (>1×10^8^ photons/second), we systemically injected 1×10^6^ CD2^KO^ CART2 cells. Retreated mice were able to sustain remission through 50 days after rescue (**Fig. 3E**), with durable responses correlating also with CD2^KO^ CART2 persistence within the peripheral blood (**Fig. 3F**). Taken together, these data suggest CD2^KO^ CART2 is active in preclinical patient-derived models of T-cell neoplasms.

### CD2 deletion decreases the anti-tumor function of CAR T-cells

While CD2^KO^ CART2 demonstrated strong activity in T-cell leukemia and lymphoma models, direct comparison with CD2^WT^ CART2 was precluded by fratricide and manufacturing failure. We therefore used anti-CD19 CAR T cells as a surrogate to assess the role of CD2 in CAR T-cell function. CD2-deficient CART19-BBz expanded comparably to CD2^WT^ during manufacturing (**Fig. S4A**) but displayed reduced cytotoxicity (**Fig. 4A, S4B**) and impaired proliferation (**Fig. S4C**) against Nalm6. In contrast, we found no significant differences in mitochondrial respiration of CD19- and CD58-activated CD2^KO^ and CD2^WT^ CART19 (**Fig. S4D**), suggesting CD2 signaling does not influence CAR T-cell metabolism.

**FIGURE 4.**
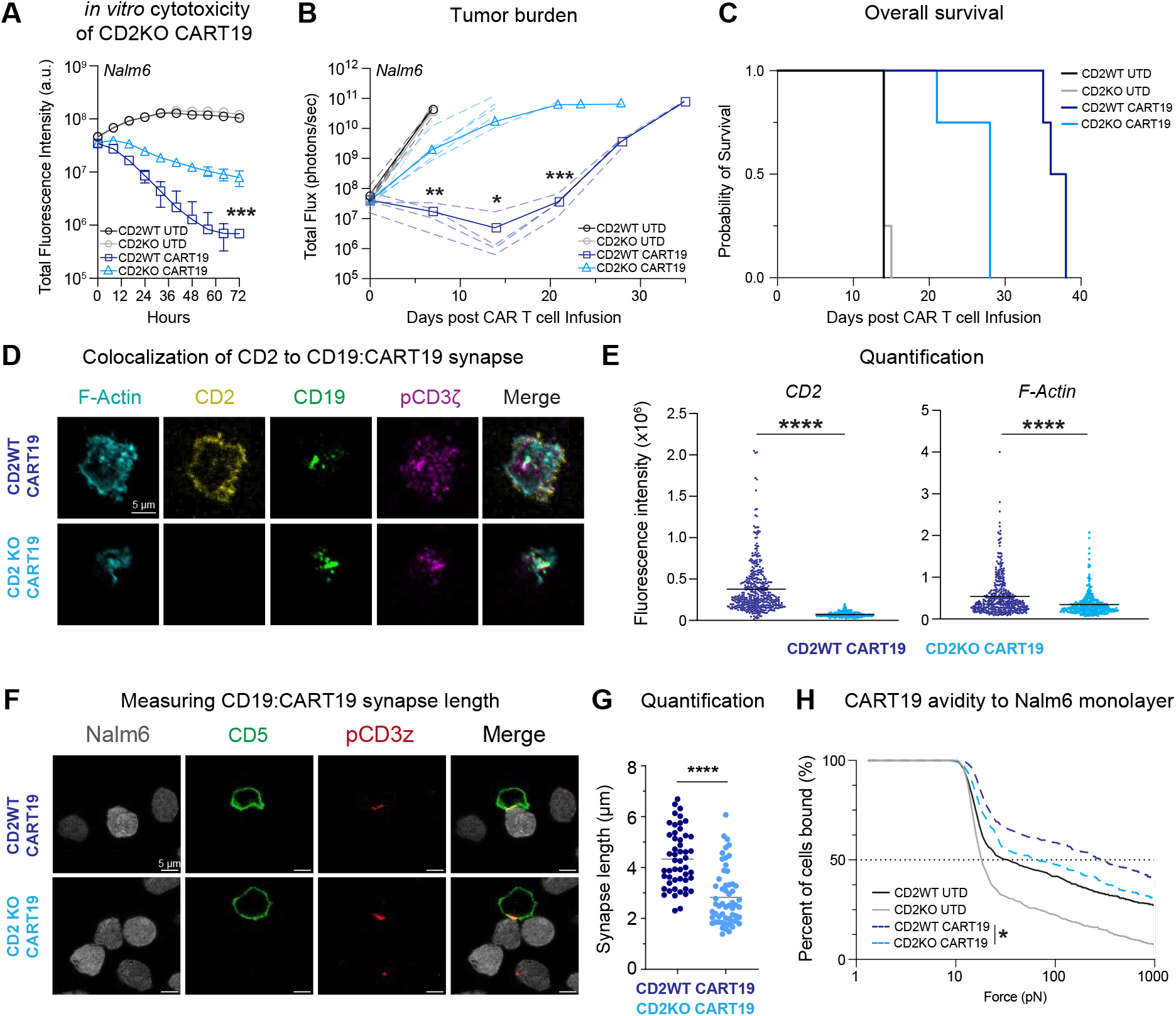
CD2 is required for optimal CAR T-cell efficacy. **(A)** Total fluorescence intensity of CD2^WT^ UTD, CD2^KO^ UTD, CD2^WT^ CART19, or CD2^KO^ CART19 cells against Nalm6 after 72 hours as continuously measured using a CellCyte Live Cell Analyzer (E:T ratio = 0.125:1). Significance determined by ANOVA with Tukey post-hoc multiple comparisons test; *** p < 0.001 relative to CD2^KO^ CART19. **(B)** Total BLI of each individual mouse treated post-T-cell infusion is shown. *NSG* mice were intravenously engrafted with 1.0 x 10^6^ Nalm6 and subsequently treated with 1.0 x 10^6^ CD2^WT^ UTD, CD2^KO^ UTD cells, 1.0 x 10^6^ CAR-positive CD2^WT^ CART19, or 1.0 x 10^6^ CAR-positive CD2^KO^ CART19 cells seven days after tumor injection. Solid lines represent the median of each cohort. Mice were monitored either until death due to any cause or euthanized at the predetermined humane endpoint (Total Flux = 10^11^ photons/sec). Significance determined by paired t-test between CD2^KO^ CART19 and CD2^WT^ CART19 at each time point; ***p < 0.001, **p < 0.01, *p < 0.05. **(C)** Overall survival of composite *NSG* mice treated with CD2^WT^ UTD, CD2^KO^ UTD, CD2^WT^ CART19, or CD2^KO^ CART19 cells (mOS: CD2^WT^ UTD: 14 days, CD2^KO^ UTD: 14 days, CD2^KO^ CART19: 28 days, CD2^WT^ CART19: 38 days). Wilcox rank-sum test was performed for all survival curve comparison between CD2^WT^ CART19 and CD2^KO^ CART19; *p < 0.05. **(D)** Representative images of immune synapses formed by CD2^WT^ CART19 *(top)* and CD2^KO^ CART19 cells *(bottom)*, visualized by planar lipid bilayer immunofluorescence microscopy against biotinylated CD19. pCD3ζ (magenta) marks CAR molecules and colocalizes with CD19 (green), delineating the CART19 synapse architecture. The experiment is designed to study the immune synapse from a top-down perspective, and what is shown is the imagine footprint of a single CAR-T cell engaging the bilayer. **(E)** Quantification of CD2 and F-actin recruitment to the CD19:CART19 immune synapses. Each data point represents an individual synapse. Significance was assessed by Student’s t-test; ****p < 0.0001. **(F)** Representative images of immunological synapses formed by CD2^WT^ CART19 (*top*) and CD2^KO^ CART19 (*bottom*), stained for, in order: NALM6 GFP, CD5, pCD3ζ, and the composite overlay. Images were acquired using a Leica Stellaris confocal microscope with a 63x/1.4 oil immersion objective. **(G)** Quantification of pCD3ζ signal length in immune synapses showed in (F). Each data point represents an individual synapse. Statistical analysis was performed using a Mann–Whitney test; ****p < 0.0001. **(H)** Quantification of CD2^WT^ UTD, CD2^KO^ UTD, CD2^WT^ CART19, and CD2^KO^ CART19 binding avidity to Nalm6 after a 15-minute co-culture using the LUMICKS z-Movi Cell Avidity Analyzer, one representative run shown. Upon application of a 1000 pN acoustic force, CD2^WT^ CAR19 T cells exhibited greater avidity toward target cells compared to CD2^KO^ CAR19 T cells, as determined by Student’s t-test (p < 0.05).

We hypothesized that CD2 loss would exert a more pronounced impact *in vivo*. Indeed, CD2^KO^ CART19-treated mice failed to adequately control Nalm6 progression compared with CD2^WT^ (**Fig. 4B**), resulting in shorter survival (**Fig. 4C**). These findings were confirmed in a different experiment with an independent donor (**Fig. S4E**). In both experiments, death was attributable, in all cases, to leukemic relapse and not graft-versus-host disease (**Fig. S4F**). Analysis of serum collected on day 10 post-infusion revealed reduced effector cytokines (i.e., IFN-γ, IL-2, sILR2α, IL-5, CXCL10/IP) in CD2^KO^ CART19 mice (**Fig. S4G**). Since CD2 signaling is known to synergize with CD28 to support TCR-dependent cytotoxic activity in endogenous T-cells(35,36), we also tested CD28-costimulated CART19 with or without CD2 in *NSG* mice engrafted with Nalm6. Again, CD2^KO^ CART19 underperformed relative to CD2^WT^ in both leukemia control and overall survival (**Fig. S4H**).

### Lack of CD2 on CAR T-cells leads to reduced T-cell activation and decreased effector function

Having established that CD2^KO^ compromises CAR T-cell activity, we next investigated the underlying mechanism. Using confocal immunofluorescence in a CD19-lipid bilayer system, we observed CD2 recruitment to the CD19:CART19 immune synapse (IS) in CD2^WT^ cells (**Fig. 4D**). Although CD2 absence did not prevent CD19:CART19 engagement, it markedly reduced F-actin accumulation, a cytoskeletal element essential for TCR-driven activation(28,52–54) (**Fig. 4E**). Accordingly, quantification of phosphorylated-CD3ζ signal length by confocal microscopy demonstrated that CD2^WT^ CART19 generated longer and more stable synapses than CD2^KO^ counterparts (**Fig. 4F, 4G**). We next evaluated whether this structural impairment translated into altered CAR T:tumor cell avidity using the Z-Movi platform. As expected, CD2 loss resulted in a significant reduction of CAR T:tumor cell binding strength (**Fig. 4H**). Collectively, these data indicate that CD2 contributes to MHC-independent IS assembly, thereby ensuring optimal CAR T-cell engagement with tumor targets.

To characterize the impact of defective immunological synapse formation in the absence of CD2, we performed single-cell RNA sequencing (scRNA-seq) on human CAR T cells from an in vivo lymphoma model. Briefly, mice were engrafted subcutaneously with OCI-Ly18 and infused with a curative dose of CD2^WT^ or CD2^KO^ CART19 (**Fig. 5A, 5B, S5A**). On day 16 post-infusion, peripheral blood mononuclear cells (PBMCs) were harvested, sample-barcoded to preserve mouse identity, and positively selected for human T-cell markers (CD2, CD3, CD5). Quality control confirmed representation from each mouse across clusters (**Fig. S5B, S5C**). Using key marker genes, we identified eight distinct T-cell subsets with comparable CD2^WT^/CD2^KO^ distributions among the clusters (**Fig. 5C, 5D, S5D**). Bulk analysis revealed reduced expression of genes critical for maintaining T cell memory in CD2^KO^ CART19, including *IL7R*(55), *TCF7* (TCF-1)(56)and *LEF1*(57)as well as decreased expression of T cell signaling molecules such as *CARD11* and *PIK3CD* (PI3Kδ). Conversely, CD2^KO^ CART19 cells had significantly increased expression of canonical exhaustion markers (i.e., *PDCD1* (*PD-1*), *HAVCR2* (*TIM-3*), *CTLA4, TIGIT, LAG3*) (**Fig. 5E**). In particular, within the GZMB^+^PRF1^+^ cytotoxic T cell subset, CD2^KO^ CART19 upregulated inhibitory regulators (*RGS16, PKMYT1*) and downregulated key components of the CD2-dependent signaling pathways, such as PI3K-Akt (*PIK3R5, AKT3, CARD11*) and MAPK pathways, including MAPK4 (**Fig. 5F**). Pathway analysis of differentially expressed genes within the CD2^KO^ cytotoxic CD8+ subset corroborated broad attenuation of T-cell activation programs and, notably, cell-to-cell adhesion in the absence of CD2 (**Fig. 5G**). Overall, these data indicate that CD2-dependent signaling supports optimal CAR T-cell activation and the maintenance of tumor-reactive phenotypes.

**FIGURE 5.**
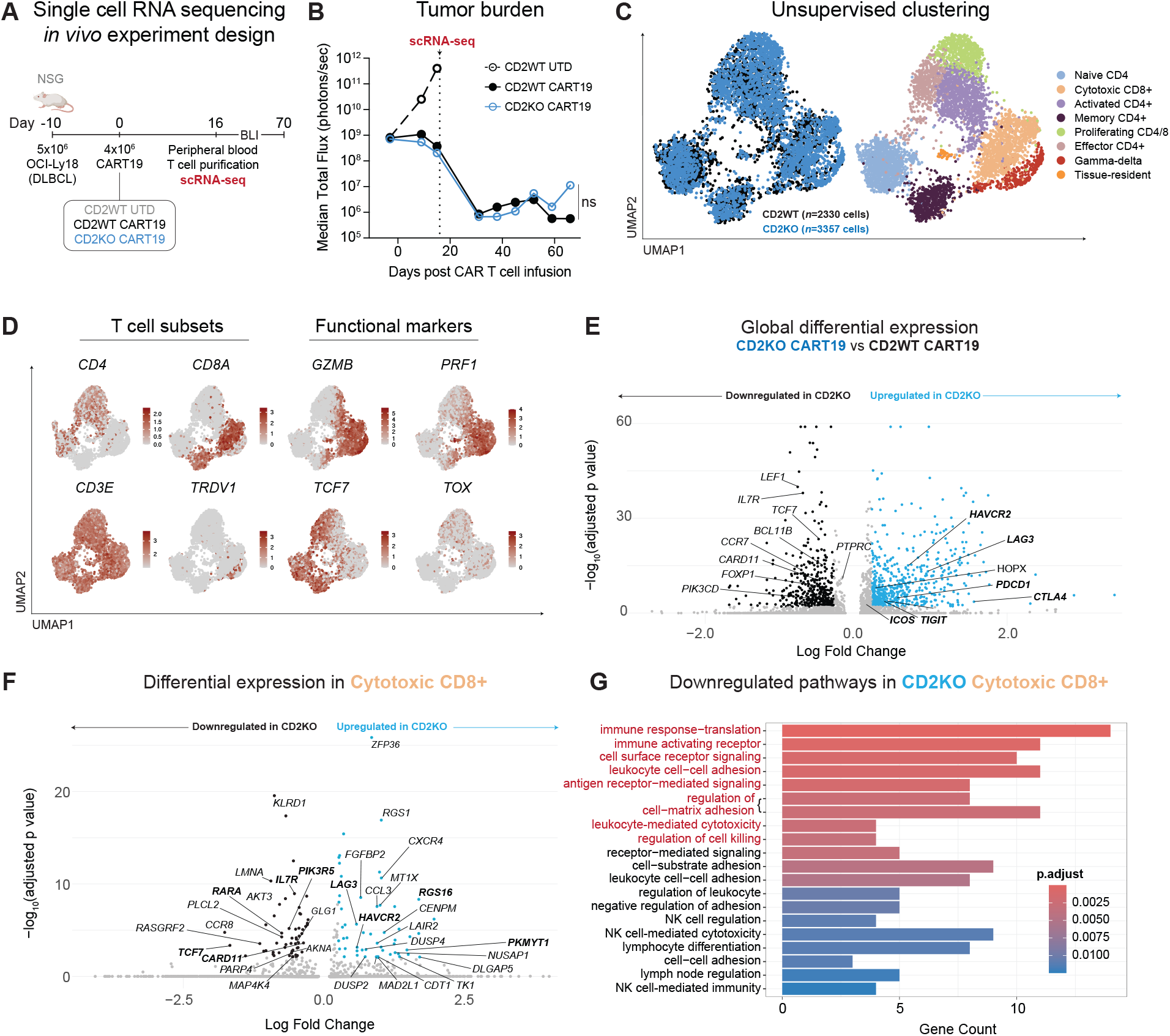
Single-cell RNA sequencing analysis of in vivo-derived CD2^KO^ CART19. **(A)** NSG mice were subcutaneously engrafted with 5.0 × 10^6^ OCI-Ly18 (diffuse large B-cell lymphoma) cells on day -10, followed by intravenous injection of 4.0 × 10^6^ Mock or CD2^KO^ CART19 cells on day 0. **(B)** Median of the bioluminescence imaging of tumor burden in NSG mice (n = 8 for CD2^WT^ CART19, n = 6 for CD2^KO^ CART19) engrafted with OCI-Ly18 cells. **(C)** Unsupervised clustering using UMAP of all cells, grouped by experimental condition (*left*) or by T cell subset as defined via the Seurat pipeline (*right*). **(D)** Expression of genes defining broad T cell subsets (CD4, CD8A, CD3E, TRDV1 (T cell receptor delta variable 1)) and other key functional T cell genes (TCF7 (Transcription Factor 7), TOX (Thymocyte Selection-Associated High Mobility Group Box), GZMB (Granzyme B), PRF1 (Perforin 1)). **(E)** Volcano plot showing significantly differentially expressed genes between CD2KO CART19 and CD2^WT^ CART19 in the bulk T cell population. **(F)** Volcano plot showing significantly differentially expressed genes between CD2^KO^ CART19 and CD2^WT^ CART19 within the Cytotoxic CD8+ T cell subset. **(G)** Pathway analysis of differentially regulated pathways in the Cytotoxic CD8^+^ T cell cluster, comparing CD2^KO^ CART19 to CD2^WT^ CART19.

### CD58 plays a critical role in CD2-mediated CAR T-cell stimulation and anti-tumor effects

Having demonstrated that CD2 signaling is critical for CAR T-cell efficacy, we next examined CD58, the predominant human CD2 ligand. Notably, CD58 is broadly expressed in PTCL, T-ALL (**Fig. S6A & S6B**)(49,58), and across diverse hematopoietic and non-hematopoietic malignancies(59). To test whether CD58 loss confers resistance to CAR T cells, we interrogated our unbiased genome-wide CRISPR-Cas9 knock-out screen of Nalm6 co-cultured with CART19, a loss-of-function platform to identify tumor-intrinsic regulators of resistance(60). This analysis revealed significant enrichment of a CD58-targeting gRNA, indicating that CD58 disruption correlates with CART19 resistance (**Fig. 6A**). To validate these findings clinically, we performed bulk RNA-sequencing on diagnostic biopsies from 17 patients with diffuse large B-cell lymphoma (DLBCL) prior to CART19 treatment (NCT02030834). Patients in the highest CD58 tertile had superior overall survival (mOS: 26.2 months) compared with those in the lowest tertiles (7.9 months) (**Fig. 6B**). Thus, tumor-associated CD58 aberrancies were hypothesized to influence CART19 responses.

**FIGURE 6.**
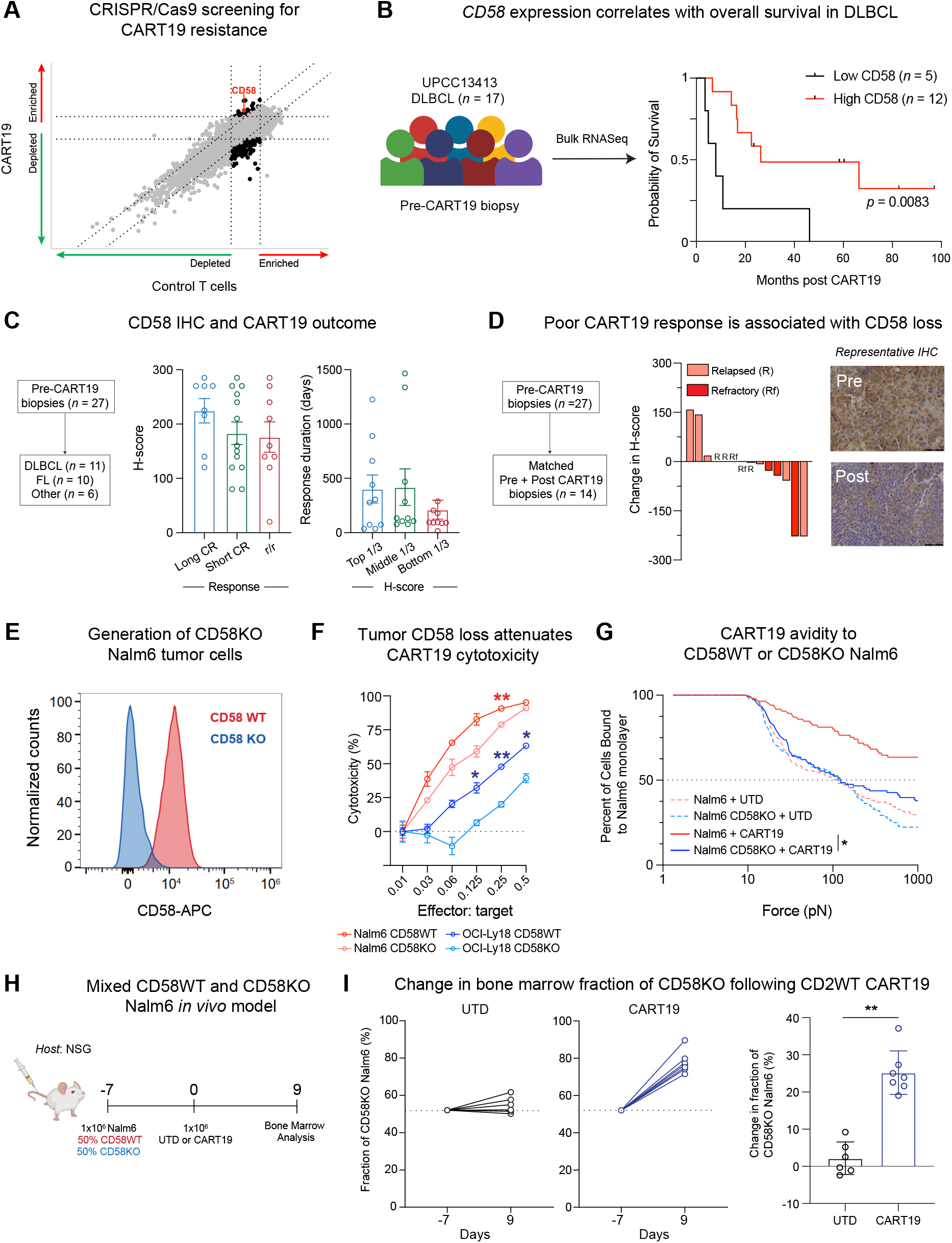
CD2:CD58 signaling axis influences CAR T-cell response. **(A)** Scatterplot of normalized MAGeCK beta scores representing enriched and depleted sgRNAs utilized in a genome-wide knock-out screen of Brunello library-edited Nalm6 cells and CART19, as previously described by Singh N, *et al*.(60). **(B)** Overall survival of 17 patients with diffuse large B-cell lymphoma (DLBCL) enrolled to UPCC13413 (NCT02030834) and stratified based on *CD58* expression, as determined by bulk tumor RNA sequencing collected prior to CART19 administration. Low CD58: Lowest 1/3 of patients based on *CD58* gene expression; High CD58: Upper 2/3 of patients based on *CD58* gene expression. **(C)** *Left:* Schematic representation of pre-CART19 biopsies from patients with aggressive B-cell lymphomas (Others: marginal zone lymphoma, grey zone lymphoma, and primary mediastinal B-cell lymphoma). *Center:* CD58 immunohistochemistry H-scores of B-cell lymphoma biopsies obtained prior to CART19 treatment. Groups include: Long CR: patients who achieved a complete remission (CR) lasting more than 1 year; Short CR: patients with a CR lasting less than 1 year; and Refractory/Relapsed: patients who experienced disease progression or relapse within 90 days of CART19 infusion. *Right:* Duration of complete response in days stratified by tertiles of pre-CART19 biopsy CD58 H-score. **(D)** *Left:* Schematic representation of the pre-and post-CART19 biopsies analyzed. *Center:* Change in CD58 H-score (*delta* H-score) in patients with available paired pre- and post-treatment biopsies. “Relapsed” (R) indicates disease progression following a documented complete response, and “Refractory” (Rf) refers to failure to achieve a complete response. *Right:* Representative CD58 immunohistochemistry images from a patient with DLBCL who experienced relapse after CART19 treatment, showing staining in the pre-treatment biopsy *(inset, top)* and the post-relapse biopsy *(inset, bottom)*. **(E)** Validation of CD58 knockout in NALM6 cells by flow cytometry. Histogram overlay showing surface CD58 expression in wild-type and CD58 knockout NALM6 cells, as assessed by flow cytometry. Loss of CD58 expression confirms effective gene disruption using the gRNA sequence listed in Table S1. **(F)** Percent cytotoxicity of CART19 cells against OCI-Ly18 and Nalm6 cells with (CD58^WT^) and without (CD58^KO^) CD58 expression across a range of effector-to-target cell ratios after 48 h coculture. Groups compared using ANOVA with Tukey post-hoc multiple comparisons test relative to CD58^WT^ for each cell type; ** p < 0.01, * p < 0.05. **(G)** Quantification of UTD or CART19 binding avidity to CD58^WT^ and CD58^KO^ Nalm6 after 15 minutes co-culture on the LUMICKS z-Movi Cell Avidity Analyzer. Upon application of a 1000 pN acoustic force, CAR19 T cells exhibited greater avidity toward NALM6 CD58^WT^ cells compared to NALM6 CD58^KO^ cells, as determined by Student’s t-test (p < 0.05). **(H)** In vivo schema to evaluate the influence of CD58 on CART19 response. Mice were co-engrafted on day –7 with a 1:1 mixture of NALM6 CD58^WT^ and CD58^KO^ cells engineered to express GFP and luciferase. On day 0, mice received 1 × 10^6^ CD2^WT^ UTD or CD2^WT^ CART19 cells. Mice were euthanized on day 9 to assess the composition of CD58^WT^ and CD58^KO^ tumor cells in the bone marrow by flow cytometry. **(I)** *Left:* Fraction of CD58^KO^ NALM6 cells in the bone marrow at day –7 and day 9 in mice treated with CD2^WT^ UTD or CD2^WT^ CART19, as assessed by flow cytometry. *Right*: Change in the fraction of CD58^KO^ cells between day –7 and day 9. Statistical significance was determined using Student’s t-test; **p < 0.01.

To gain further clinical insight, we assessed biopsies from r/r B-cell lymphomas obtained before commercial CART19 (n=27) or after relapse (n=14) at the University of Pennsylvania. Pre-treatment CD58 H-score showed a trend toward higher levels in patients achieving long complete responses (CR >1 year; 224.5±21.1) compared with short CR (<1 year; 183.0±20.0) or non-responders (176.1±28.0). Notably, pre-CART19 biopsies in the lowest H-score tertile also showed a trend toward shorter response duration (**Fig. 6C**). Furthermore, in patients with paired biopsies available, CD58 Immunohistochemistry expression was reduced at relapse in 50% (7/14) of cases (**Fig. 6D**). This trend was independent of B-cell lymphoma subtype or commercial CAR T-cell product infused (data not shown).

Lastly, we tested whether CD58 deletion on tumor cells would attenuate CART19 activity. After generating CD58-deficient Nalm6 (**Fig. 6E**) and OCI-Ly18 cell lines, we observed reduced CART19-mediated cytotoxicity in the CD58^KO^ setting (**Fig. 6F**). Mechanistically, CD58 deletion impaired CART19 avidity, mirroring the phenotype seen with CD2 loss (**Fig. 6G**). To test whether CD58 loss conferred a growth advantage under CAR T-cell pressure, *NSG* mice were engrafted with a 1:1 mixture of CD58^WT^ and CD58^KO^ Nalm6 cells and treated with either UTD or CART19 (**Fig. 6H**). In the absence of CART19, CD58^KO^ cells showed no proliferative advantage over CD58^WT^ counterparts. In contrast, CART19 treatment induced selective enrichment of CD58^KO^ cells, with ∼25% increase over baseline **(Fig. 6I)**. Collectively, these data demonstrate that CD58 loss promotes immune evasion under CAR T-cell pressure.

### A PD-1:CD2 switch receptor rescues the lack of CD2 engagement on CART19

Having demonstrated that the CD2:CD58 axis is critical for CART efficacy, we explored translational strategies to bypass the absence of CD2 signaling and preserve synapse formation, co-stimulation, and anti-tumor activity. As a first approach, we engineered a third-generation CART19 incorporating the intracellular CD2 signaling domain (CART19-iCD2) (**Fig. S7A**). While this design enhanced *in vitro* cytotoxicity (**Fig. S7B**), it failed to reproduce this effect *in vivo* against CD58^KO^ Nalm6 (**Fig. S7C**).

We hypothesized that a novel *in trans* switch receptor with a membrane-proximal intracellular CD2 domain could restore CD2 signaling in CAR T cells lacking CD2:CD58 engagement (**Fig. 7A**). PD-1 was selected as the extracellular domain due to its proximity to CD2 in T-cell synapse(30) and the consistent expression of PD-L1 in PTCL, T-ALL(49,58) (**Fig. S6A, S6B**), B-cell malignancies(61) and solid cancer(62). Thus, a PD-1:CD2 receptor would exploit PD-L1 on tumor cells to trigger CD2 signaling alongside CD19:CART19 engagement. Using a P2A lentiviral vector (**Fig. S7D**), we achieved high co-expression of PD-1:CD2 with CART19 (**Fig. 7B, Fig. 7C**), while preserving manufacturing kinetics (**Fig. S7E**). To test the PD-1:CD2 functionality, CART cells were stimulated on plates coated simultaneously with CD19, CD58, and PD-L1; cell lysates were analyzed by immunoblot for phospho-Lck, a key CD2 signaling readout(63) (**Fig. 7D**). CD2^KO^ CART19 showed reduced p-Lck compared to CD2^WT^, whereas CART19 cells expressing the full-length PD-1:CD2 receptor restored p-Lck, in contrast to the truncated PD-1 receptor lacking the intracellular CD2 domain (**Fig. 7E**). The PD-1:CD2 receptor also enhanced killing of CD58-deficient Nalm6 expressing different levels of PD-L1, demonstrating activity even under limited PD-L1 ligand availability (**Fig. S7F**). Finally, using a Jurkat reporter for NFAT, NFκB, and AP-1, co-expression of the full-length PD-1:CD2 switch receptor with CAR19 markedly increased transcriptional activity upon co-culture with either CD58^WT^ or CD58^KO^ Nalm6-PD-L1+ cells. This effect was absent in the truncated receptor, confirming that intracellular CD2 signaling specifically mediates transcriptional reprogramming (**Fig. S7G, S7H, S7I**). Overall, these results establish that the PD-1:CD2 switch receptor restores CAR T-cell function via selective activation of intracellular CD2 signaling.

**FIGURE 7.**
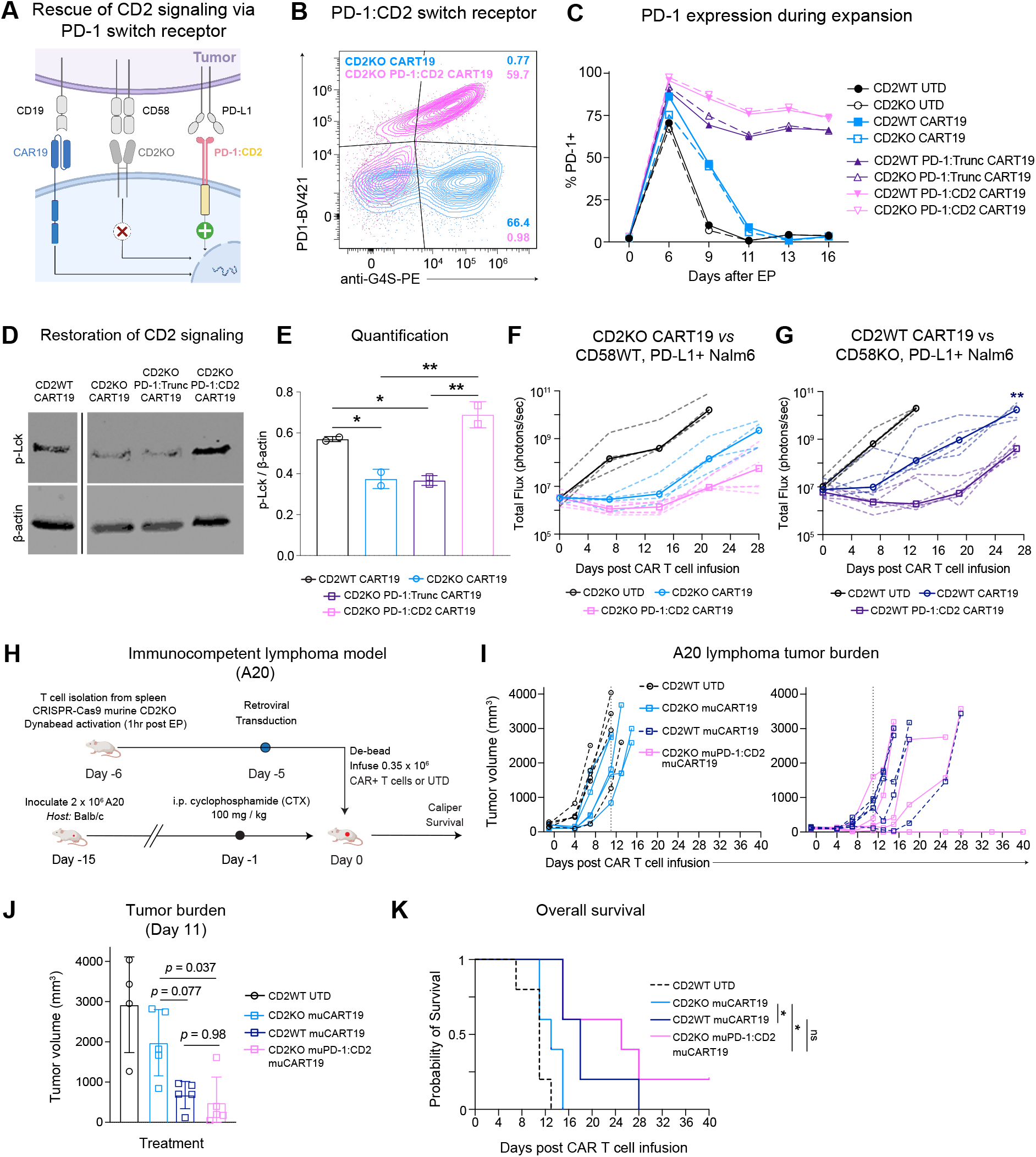
Functional and in vivo validation of the PD-1:CD2 switch receptor in CART19 cells. **(A)** Schema of a PD-1:CD2 switch receptor in CD2^KO^ CART19. **(B)** Representative flow cytometry immunophenotyping on day 16 of T-cell expansion of CD2^KO^ CART19 and CD2^KO^ PD-1:CD2 CART19; *y-axis:* PD-1–BV421 (representing the extracellular domain of the PD-1:CD2 switch receptor) and *x-axis:* G4S–PE (CAR19). **(C)** PD-1 expression over the course of T-cell expansion, representing either the full PD-1:CD2 switch receptor, a truncated version lacking the intracellular CD2 domain, or endogenous physiological PD-1. **(D)** Immunoblot of p-Lck protein in CD2^WT^ CART19, CD2^KO^ CART19, and CD2^KO^ CART19 expressing truncated or full PD-1:CD2 switch receptors following 20 minutes of stimulation on plates simultaneously coated with CD19, PD-L1, and CD58. Bands were derived from the same gel; non-contiguous lanes were juxtaposed, and black lines indicate where the gel was cut. No other modifications were made. **(E)** Densitometry quantification of p-Lck normalized to β-actin from the protein lysates shown in Figure 7D. Statistical analysis was performed by one-way ANOVA followed by Tukey’s post-hoc multiple comparisons test. **p < 0.01, *p < 0.05. **(F)** In vivo experiment where NSG mice were intravenously injected with 1.0 x 10^6^ PD-L1-overexpressing CD58^WT^ Nalm6 and subsequently treated with 1.0 x 10^6^ CD2^KO^ UTD (n=4), CD2^KO^ CART19 (n=5), or CD2^KO^ CART19 expressing the PD-1:CD2 switch receptor (n=6) cells 7 days after leukemia injection (day 0). Total BLI of individual mice treated (dashed lines). Solid lines represent the median of each cohort. Groups were compared using ANOVA with Tukey post-hoc multiple comparisons at each time point measurement, **p < 0.01, * p < 0.05. **(G)** In vivo experiment where NSG mice were intravenously injected with 1.0 x 106 PD-L1-overexpressing CD58^KO^ Nalm6 and subsequently treated with 1.0 x 106 CD2^WT^ UTD (n=4), CD2^WT^ CART19 (n=5), or CD2^WT^ CART19 expressing the PD-1:CD2 switch receptor (n=6) cells 7 days after leukemia injection (day 0). Total BLI of individual mice treated (dashed lines). Solid lines represent the median of each cohort. Groups were compared using ANOVA with Tukey post-hoc multiple comparisons at each time point measurement. **(H)** Schematic of the A20 lymphoma model (see Methods). On day 0, 1 × 10^6^ CD2^WT^ UTD, CD2^WT^ muCAR19, CD2^KO^ muCAR19, or CD2^KO^ muCAR19 expressing the muPD-1:CD2 switch receptor were infused intravenously. **(I)** A20 tumor growth curves showing individual mouse replicates following infusion of murine UTD or CART19 T cells. *Left*: CD2^WT^ UTD and CD2^KO^ muCAR19. *Right*: CD2^WT^ muCAR19 and CD2^KO^ muCAR19 expressing the muPD-1:CD2 switch receptor. The vertical dashed line indicates the Day 11 measurement. Tumor volume was calculated using the formula: (length × width^2^) /2. **(J)** Tumor volume at Day 11 post-infusion. Statistical analysis was performed by one-way ANOVA followed by Tukey’s post-hoc multiple comparisons test. **(K)** Overall survival curves, Wilcox rank-sum test was performed for all survival curve comparisons between CD2^KO^ muCART19 vs CD2^WT^ muCART19, CD2^KO^ muCART19 vs CD2^KO^ muCART19 expressing the muPD-1:CD2 switch receptor, and CD2^WT^ muCART19 vs CD2^KO^ muCART19 expressing the muPD-1:CD2 switch receptor. * p < 0.05.

Lastly, to evaluate the *in vivo* efficacy of the PD-1:CD2 switch receptor, *NSG* mice were engrafted with PD-L1^+^ Nalm6 and treated with CD2^KO^ CART19 or CD2^KO^ CART19 co-expressing PD-1:CD2. Mice receiving switch receptor–modified cells exhibited superior leukemia control and significantly prolonged survival **(Fig. 7F, S7J)**. A similar rescue was observed in CD58^KO^ Nalm6, where CD2^WT^ PD-1:CD2 CART19-BBζ outperformed CD2^WT^ CART19 **(Fig. 7G, S7K)**. To extend these findings to an immunocompetent setting, we first disrupted CD2 in murine T-cells via CRISPR **(Fig. S7L, Table S2)** and then co-expressed a murine PD-1:CD2 receptor (muPD-1:CD2) with a murine anti-CD19 CAR (muCART19) using a T2A construct (**Fig. S7M**). BALB/c mice were first engrafted with A20 lymphoma, underwent CTX lymphodepletion at day -1, and then infused with murine CART19 cells **(Fig. 7H)**. In this model, CD2^KO^ muCART19 exhibited impaired tumor control compared to CD2^WT^ counterpart. Interestingly, co-expression of muPD-1:CD2 in CD2^KO^ muCART19 restored anti-tumor efficacy, leading to tumor control and survival comparable to CD2^WT^ muCART19 **(Fig. 7I, 7J, 7K)**. Collectively, these results demonstrate that defects in the CD2:CD58 axis can be overcome by PD-1:CD2–mediated signaling rescue.

## DISCUSSION

Here, we report the successful development and optimization of a CAR T-cell therapy directed against CD2 for the treatment of CD2-expressing malignancies.

Several technical and clinical obstacles have limited the success of previous T-cell-directed cellular immunotherapies. First, shared antigen expression between malignant and healthy T cells leads to fratricide. Clinical studies of unedited CART5, without concurrent CD5 deletion, showed limited persistence and suboptimal outcomes(15). These findings underscore the need to abolish fratricide when targeting pan–T-cell antigens. We found that elimination of endogenous CD2 effectively prevented CART2 fratricide while preserving T-cell activation through MHC-independent tumor recognition.

Second, antigen-negative escape following CAR T-cell treatment is common. As observed with CD19 loss after CART19 therapy, antigen downregulation or clonal selection can emerge(64–66). In early studies of CD7-directed CAR T cells, up to 30% of patients developed CD7-negative escape(27), reflecting the redundant role of CD7 in T-cell biology and its propensity for tumor downregulation(20,67). While antigen escape is possible with any single-antigen CAR, the threshold may be higher for CD2 due to its function in co-stimulation and immune synapse stabilization. To address tumor heterogeneity, we explored two strategies: targeting a uniformly expressed antigen and sequential CAR therapies. Here, we show that CD2 is highly and uniformly expressed in T-lymphoma and T-ALL, supporting its use as a stable target. Moreover, CART2 administered at relapse in patient-derived T-ALL progressing after CART5 induced complete remission in CD5-negative tumors. These results highlight CD2 as a robust antigen and support sequential targeting to overcome antigen-negative escape.

UCART2, an allogeneic “off-the-shelf” anti-CD2 CAR T product, was developed with TCR knockout to prevent graft-versus-host disease(68). In contrast, the CART2 described here is autologous, eliminating the need for TCR deletion. Preserving the TCR may sustain adaptive immunity against pathogens and mediate a graft-versus-tumor effect. Moreover, UCART2 required repeated co-administration of rh-IL7 for durable efficacy, a potential drawback in T-cell neoplasms responsive to IL-7(69). For these reasons, our strategy focuses on optimizing autologous CART2 to induce complete remission in T-ALL as a bridge to transplant, as achieving CR before allogeneic stem-cell transplantation remains the strongest prognostic factor for long-term cure(70–72).

Although we observed long-term remissions after CART2 treatment in preclinical models, we also established that CD2 loss diminishes CAR T-cell activity using CART19 as a surrogate. Deletion of CD2 in CAR T cells or CD58 in tumor cells impaired responses by disrupting immune synapse organization, reducing avidity, and limiting CD2-mediated co-stimulation. Interestingly, our finding was recently corroborated by Zhu et al., who demonstrated that CD2 expression levels correlate with CAR T-cell immunological synapse quality (73). Clinically, our data suggest that CD58 expression may serve as a predictor of CART19 efficacy in advanced B-cell lymphomas. This is consistent with prior reports linking CD58 downregulation to poor prognosis in B-cell lymphoma(74–76). Our analysis of tisagenlecleucel-treated patients at the University of Pennsylvania, together with a cohort of 51 DLBCL patients treated at Stanford with axicabtagene ciloleucel(77), further supports CD58 immunohistochemistry as a potential biomarker of CART19 activity. Larger prospective studies are warranted to validate CD58 as a predictive biomarker of CAR T-cell efficacy.

Finally, a key novelty of this work is the development of a strategy to overcome loss of CD2:CD58 signaling. We restored CD2 activity in CAR T cells using a synthetically engineered PD-1:CD2 switch receptor, effective in both immunodeficient and immunocompetent models. This receptor rescued in vivo CAR T-cell efficacy more efficiently than intracellular CD2 insertion in cis with CART19, likely due to the membrane-proximal positioning of the CD2 domain enhancing costimulation upon PD-1–PD-L1 engagement. Notably, loss of CD58 expression correlated with increased PD-L1 on tumor cells(78). Thus, the PD-1:CD2 switch receptor is expected to function in diverse contexts where the CD2:CD58 axis is disrupted, including CD2 deletion in CAR T cells, absence of CD58 in the tumor, or altered CD58 expression.

In summary, this study identifies human CD2 as a critical regulator of CAR T-cell activity. Using primary patient samples *in vitro* and *in vivo*, we establish CD2 as a promising target for cellular therapies against CD2-expressing neoplasms. While disruption of the CD2:CD58 axis attenuates CAR T-cell function, these defects can be rescued by intracellular CD2 signaling. Our findings support the clinical development of CART2 in T-cell neoplasms and the design of CD2-based strategies to enhance adoptive T-cell therapies.

## SUPPLEMENTAL FIGURE LEGENDS

**SUPPLEMENTAL FIGURE 1. Individual T-ALL patient surface antigen and gene expression. (A)** Normalized surface antigen expression profiles from 40 diagnostic T-ALL samples obtained from patients enrolled in the phase 3 clinical trial AALL0434, analyzed by CITE-seq. **(B)** Surface antigen and gene expression in developing thymocytes. Antibody-derived-tag (ADT) surface and mRNA expression for CD2, CD3, CD5, CD7, and CD4/8A from n=16,119 healthy-donor thymocytes profiled by CITE-seq. Cell types are organized from left to right in order of T-cell development; thymic B-cells are shown as a relative control. Cycling CD4/8+ double-positive (DP) thymocytes are represented by “DP(s)” and mature, CD4 or CD8 positive, lineage-restricted α/β (CCR7+, CCR9-) T-cells are indicated by “α/β(m)”.

**SUPPLEMENTAL FIGURE 2. Engineering and optimization of anti-CD2 CAR T-cells. (A)** *Left:* CD2 knock-out efficiency of three CD2-specific gRNAs (CD2gRNA08, CD2gRNA09, and CD2gRNA10) following electroporation (EP) in Jurkat cells. Mock curve represents EP alone without CD2-specific guide RNA to serve as a negative control. KO efficiency was quantified based on total percentage of viable Jurkat cells lacking CD2 expression as assessed by flow cytometry. One donor-paired electroporation course is shown. *Right:* Representative flow cytometry histograms of CD2 expression five days following EP with CD2gRNA8. Dashed curve: Isotype-APC antibody. **(B)** *Left*: Representative flow cytometry histograms of anti-CD2 CAR extracellular expression at the end of expansion and following one freeze/thaw cycle. *Right*: Transduction efficiency of the top three highest expressed anti-CD2 CAR constructs (CAR #1, CAR #2, CAR #6) at the end of CD2^KO^ CART2 manufacturing. Each data point represents a unique, healthy T-cell donor. UTD: untransduced. **(C)** CD4^+^ and CD8^+^ T-cell memory phenotypes of Mock UTD, CD2^KO^ UTD, and CART2 at the end of expansion. T_naive_, naïve T-cells, CD45RA^+^CCR7^+^; T_CM_, central memory T-cells, CD45RA^-^CCR7^+^; T_EM_, effector memory T-cells, CD45RA^-^CCR7^-^; T_EMRA_, effector memory T-cells re-expressing CD45RA, CD45RA^+^CCR7^-^; n= 2 donors. **(D)** Total percentage of CD8^+^ T-cells with expression of IL-2 (*left*), TNFα (*middle*), and IFN-γ (*right*) in Mock UTD, CD2^KO^ UTD, and CART2. T-cells were stimulated with R10 (unstimulated, green), PMA/Ionomycin (red), CD3/CD28 Dynabeads (blue), or cocultured with Jurkat cells (yellow) for 24 hours; n=2 technical replicates. **(E)** T cell proliferation of CD2^WT^ UTD, CD2^KO^ UTD, and CD2^KO^ CART2 cells co-cultured with irradiated Jurkat cells. T cells were labeled with CellTrace Violet prior to co-culture and analyzed by flow cytometry after 5 days. **(F)** T cell proliferation of CD2^WT^ UTD, CD2^KO^ UTD, and CD2^KO^ CAR2 T cells stimulated with CD3:CD28 Dynabeads at a 2:1 bead-to-cell ratio. T cells were labeled with CellTrace Violet prior to stimulation and analyzed by flow cytometry after 5 days. **(G)** Percent cytotoxicity of UTD, CART19, or CD2^KO^ CART2 cells against CD19+ B-cell acute lymphoblastic leukemia cells (Nalm6) after 72 hours co-culture (effector:target (E:T) ratio = 0.25:1).

**SUPPLEMENTAL FIGURE 3. CART2 *in vivo* validation against primary T-ALL. (A)** Schema of *PDX* T-ALL generation. T lymphoblasts from a pediatric patient enrolled in COG AALL1231 were transduced with a lentiviral dual GFP-Luciferase reporter plasmid *in vitro*. Once stably integrated, human T lymphoblasts were then serially passaged in *NSG* mice. At passage 2 (P2) human T lymphoblasts were collected from mouse peripheral blood, validated for CD2 expression by flow cytometry, and subsequently used for engraftment. Patient clone TH20 was used for all subsequent experiments(39). **(B)** Representative bioluminescent images (BLI) from one experiment using one independent T-cell donor for CART2 manufacturing. Notably, leukemic involvement of the central nervous system (brain) was successfully cleared by Day 14 (D14) following systemic CART2 treatment (yellow arrowheads) **(C)** Replicative bioluminescence imaging of leukemic burden (*left*) and overall survival (*right*); n= 3-7 mice per group. Wilcox rank-sum test was performed for survival curve comparisons, p=0.0004. **(D)** CD3^+^ T-cell quantification of mouse peripheral blood in viable mice at Day 28 and Day 100 post-T-cell treatment. **(E)** Body mass of each PDX T-ALL mouse in both T-cell donor cohorts. Deaths attributable to xenogenic graft-versus-host disease (arrowhead, red highlight) or disease relapse (asterisk, orange highlight) are indicated.

**SUPPLEMENTAL FIGURE 4. CD2 loss alters CART19 activity. (A)** Population doublings of two, independent, healthy T-cell donor expansions of CART19 with or without CD2 and either a 4-1BBz (CART19-BBz) or CD28z (CART19-28z) costimulatory domain. **(B)** Cytotoxicity assay after 72 hours of co-culture between CD2^WT^ UTD, CD2^KO^ UTD, CD2^WT^ CART19, or CD2^KO^ CART19 T cells and luciferase-expressing NALM6 target cells. Cytotoxic activity was measured by luminescence reduction, which reflects viable tumor cell content. Effector-to-target (E:T) ratios tested were 0.25:1, 0.125:1, and 0.0625:1. **(C)** Proliferation assay of CD2^WT^ UTD, CD2^KO^ UTD, CD2^WT^ CAR19, and CD2^KO^ CAR19 T cells co-cultured with irradiated NALM6 cells (E: T ratio of 0.25:1). T cells were labeled with CellTrace Violet prior to co-culture and analyzed by flow cytometry after 5 days. The graph shows the fold change in the absolute number of CD3^+^ T cells compared to baseline (day 0), measured using counting beads. **(D)** Metabolic profiling of CD2^WT^ UTD, CD2^WT^ CAR19, and CD2^KO^ CAR19 T cells using a *Seahorse XF* assay. 3×10^6^ CAR+ T cells (or equivalent numbers of UTD) were cultured for 48 hours on plates coated simultaneously with recombinant CD19 and CD58 proteins prior to analysis (ns, Ordinary one-way ANOVA). **(E)** Replicative bioluminescence imaging of leukemia burden engrafted with 1.0 x 10^6^ Nalm6 (*left*) and overall survival (*right*) of composite *NSG* mice treated with 1.0 x 10^6^ Mock UTD, CD2^KO^ UTD, CD2-wild type (CD2^WT^) CART19-BBz, and CD2^KO^ CART19-BBz cells from a separate, independent healthy T-cell donor. n= 3-6 mice per group. **(F)** Body mass of each NSG mouse in both donor cohorts. **(G)** Serum cytokine quantification of *NSG* mouse plasma fractionated from peripheral blood at Day 7 post CD2^WT^ UTD, CD2^KO^ UTD, CD2^WT^ CART19, and CD2^KO^ CART19 treatment, as determined by Luminex multiplex assay. **(H)** *Left:* Bioluminescence imaging (BLI) of leukemia burden in NSG mice engrafted with 1.0 × 10^6^ NALM6 cells and treated with 1.0 × 10^6^ T cells: CD2^WT^ UTD, CD2^KO^ UTD, CD2^WT^ CART19-28z, or CD2^KO^ CART19-28z. Dashed lines represent total BLI signal from individual mice over time; solid lines indicate the median signal per group. Statistical analysis was performed by one-way ANOVA followed by Tukey’s post-hoc multiple comparisons test. *Right:* Overall survival of NSG mice from the experiment shown in (H). Wilcoxon rank-sum test was used to compare CD2^WT^ CART19 and CD2^KO^ CART19 groups; **p < 0.01, *p < 0.05.

**SUPPLEMENTAL FIGURE 5. Ancillary single-cell RNA sequencing analyses of *in vivo*-derived CD2^KO^ CART19 cells. (A)** Immunophenotyping of CD2^WT^ UTD (untreated, *left*) and CD2^WT^ CART19 (*middle*) T cells prior to injection into NSG mice, corresponding to Figure 5A. To ensure complete CD2 deficiency for single-cell analyses, CD2^KO^ CART19 cells were additionally FACS-sorted based on CD2 expression prior to infusion (*right*). **(B)** Schematic of the workflow on day 16 post-infusion for mouse-specific barcoding. Following PBMC isolation, red blood cells were lysed, and cells from each mouse were individually stained for T cells (CD2/CD3/CD5-APC) and uniquely tagged with a barcoding antibody (TotalSeq-A) targeting CD45 and MHC Class I. CD2^WT^ and CD2^KO^ T cell populations were then loaded as separate samples onto the 10X Chromium Controller for single-cell RNA sequencing. **(C)** UMAP of the distribution of TotalSeq-A barcodes identifying individual mice in the Mock CART19 (n = 8) and CD2^KO^ CART19 (n = 6) groups. **(D)** Single cell expression levels of key genes involved in T cell activation (*LCK, VAV1, PLCH1)* and exhaustion (*LAG3, HAVCR2, PDCD1*) (*left)*, as well as transcription factors (*TBX21, GATA3, RORC, and FOXP3*) associated with distinct T cell subset phenotypes (*right)*.

**SUPPLEMENTAL FIGURE 6. CD58 and PD-L1 RNA expression across T-ALL and T-lymphoma subtypes. (A)** CD58 and PD-L1 RNA expression levels across T-ALL subtypes from Pölönen et al., *Nature Cancer* (2024). Subtypes include: TME-enriched, TLX1 (HOX11), ETP-like, NUP98-rearranged, TLX3 (HOX11L2), LMO2-like, TAL1 DP-like, NKX2-5, STAG2/LMO2, BCL11B-altered, MLLT10-rearranged, NUP214-rearranged, HOXA9 TCR-positive, TAL1-like, NKX2-1, KMT2A-rearranged, and SPI1-altered. (**B)** CD58 and PD-L1 RNA expression in Diffuse Large B-cell Lymphoma and T-cell lymphoma subtypes from Fiore et al., Cell Reports Medicine (2025). Subtypes include: BIA-ALCL (Breast Implant– Associated Anaplastic Large Cell Lymphoma), ALCL ALK^+^ (Anaplastic Large Cell Lymphoma, ALK-positive), ALCL ALK^−^ (Anaplastic Large Cell Lymphoma, ALK-negative), DLBCL (Diffuse Large B-Cell Lymphoma), MF (Mycosis Fungoides), AITL (Angioimmunoblastic T-cell Lymphoma), PTCL-NOS (Peripheral T-cell Lymphoma, Not Otherwise Specified), cALCL (Cutaneous Anaplastic Large Cell Lymphoma), and T-PLL (T-cell Prolymphocytic Leukemia).

**SUPPLEMENTAL FIGURE 7. CD2:CD58 signaling axis rescue strategies. (A)** Schema of *in cis* CD2 intracellular rescue in CD2-deficient CART19. **(B)** *In vitro* cytotoxicity of CD2^KO^ CART19.28z and CD2^KO^ CART19.28.CD2z (CART19-iCD2) 72 hours after coculture with either CD58^WT^ or CD58^KO^ OCI-Ly18 cells. Percent cytotoxicity for each CART19 group against CD58^KO^ OCI-Ly18 was normalized against percent cytotoxicity against CD58^WT^ OCI-Ly18. Two-way ANOVA with Tukey multiple comparison post-hoc test was used to determine statistical significance ** p < 0.01, * p < 0.05. **(C)** *NSG* mice were intravenously injected with 1.0 x 10^6^ CD58^WT^ Nalm6 and subsequently treated with 1.0 x 10^6^ untransduced T-cells (CD2^WT^ UTD, n=3), 1.0 x 10^6^ CAR-positive CD2^KO^ CART19.28z (n=6), or 1.0 x 10^6^ CAR-positive CD2^KO^ CART19-iCD2 (n=6) seven days after leukemia injection. Total BLI *(left)* and overall survival (*right*) is shown; ns = not significant. **(D)** Construct maps of a PD-1:CD2 intracellular signaling domain switch receptor integrated with a P2A self-cleaving peptide and the CAR19.BBz construct, and its corresponding truncated version lacking the intracellular CD2 domain. **(E)** Population doublings, as calculated by normalized viable cell count relative to the total number of cells initially electroporated, of each product cohort derived from a single healthy T-cell donor. **(F)** Cytotoxicity of CD2^WT^ UTD, CD2^WT^ CART19, CD2^WT^ PD-1 truncated CART19, and CD2^WT^ PD-1:CD2 CART19 against CD58^KO^ NALM6 cells with variable surface PDL1 expression. PDL1-expressing CD58^KO^ NALM6 cells were generated by lentiviral transduction with a PD-L1-overexpressing vector and subsequently mixed at defined ratios with PD-L1 wild-type CD58^KO^ NALM6 cells to obtain target populations with approximately 0%, 20%, 45%, and 100% PDL1^+^ cells. **(G)** Schematic representation of the Jurkat triple reporter cell line, engineered to report T-cell activation through NFAT-GFP, NF-κB-mCherry, and AP-1-CFP fluorescent readouts. **(H)** Fold change in MFI of NFAT-GFP, NF-κB-mCherry, and AP-1-CFP in Jurkat triple reporter cells after 24 hours of co-culture with CD58^WT^ NALM6 cells. Conditions include CD2^WT^ UTD, CD2^WT^ CART19, CD2^WT^ CART19 expressing the truncated switch receptor, and CD2^WT^ CART19 expressing the full PD-1:CD2 switch receptor (E:T 1:1). **(I)** Fold change in MFI of NFAT-GFP, NF-κB-mCherry, and AP-1-CFP in Jurkat triple reporter cells after 24 hours of co-culture with CD58^KO^ NALM6 overexpressing PD-L1. Conditions include CD2^WT^ UTD, CD2^WT^ CART19, CD2^WT^ CART19 expressing the truncated switch receptor, and CD2^WT^ CART19 expressing the full PD-1:CD2 switch receptor (E:T 1:1). **(J)** Overall survival curves relative to Fig. 7F, Wilcox rank-sum test was performed for all survival curve comparisons between CD2^KO^ CART19 and CD2^KO^ CART19 expressing the PD-1:CD2 switch receptor (p=0.001). **(K)** Overall survival curves relative to Fig. 7G, Wilcox rank-sum test was performed for all survival curve comparison between CD2^WT^ CART19 and CD2^WT^ CART19 expressing the PD-1:CD2 switch receptor (p=0.0316). **(L)** Knockout efficiency of murine CD2 measured by flow cytometry following gRNA screening. **(M)** Expression of the murine switch receptor (murineCAR19_T2A_murine_PD-1:CD2) assessed by flow cytometry in the murine CD2^KO^ T-cell population at Day 8 of murine T-cell expansion.

**Table S1.**
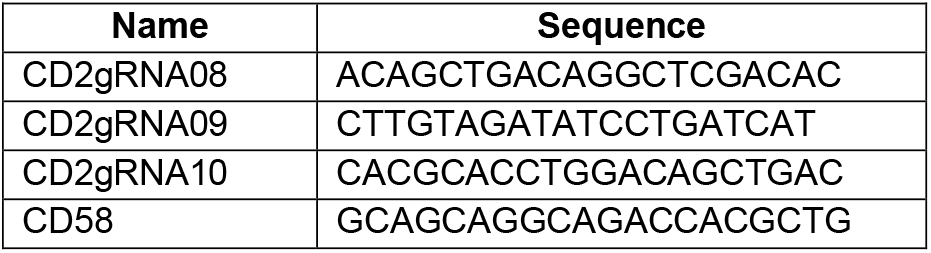
CD2 and CD58 gRNA sequences tested.

**Table S2.**
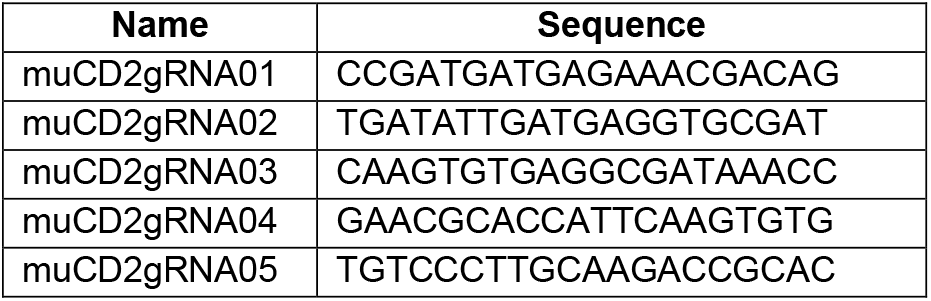
Screening of gRNAs for Murine CD2 Knockout.

**Table S3.**
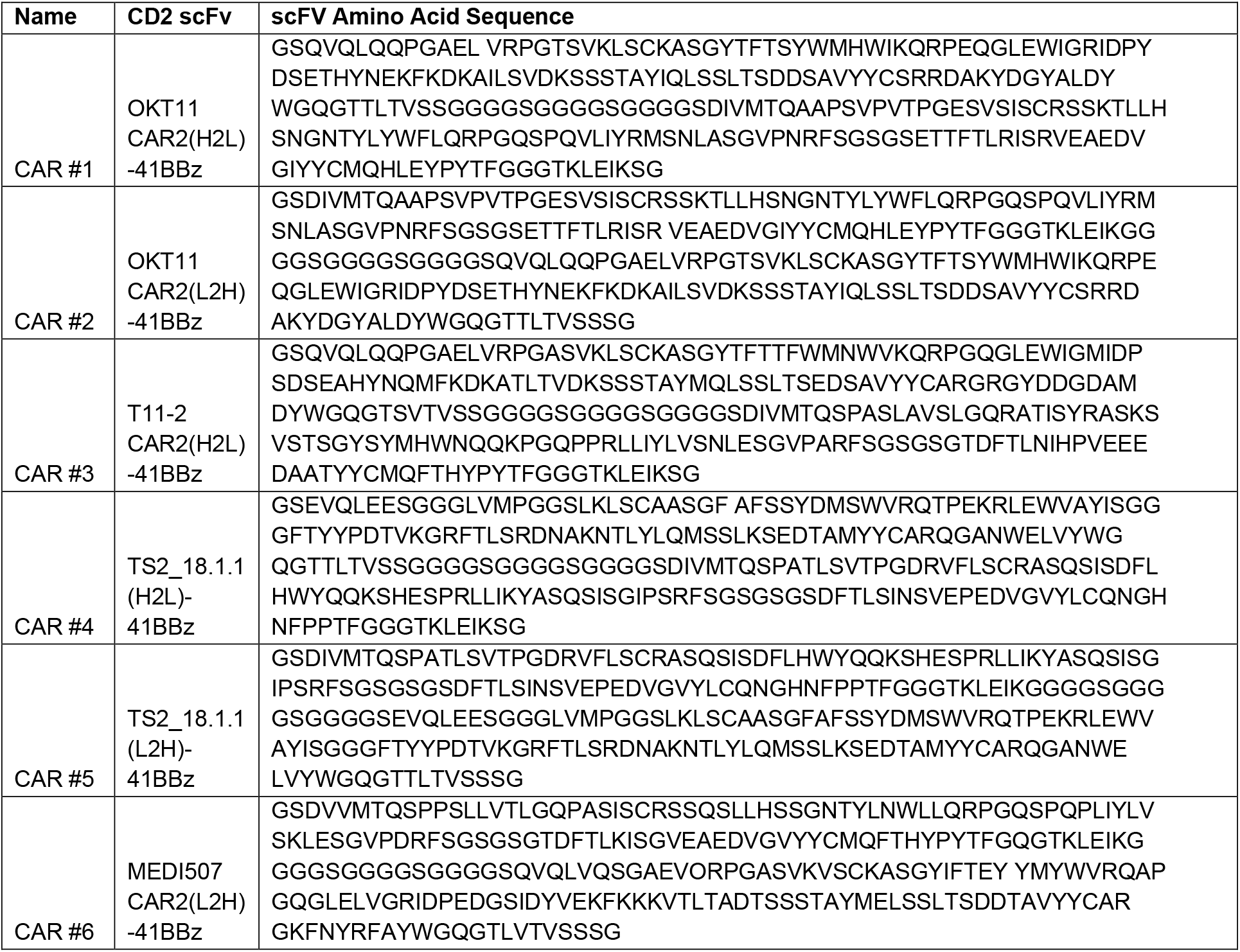
CAR2 scFv antibody clones.

## Notes

### Competing Interest Statement

A.C. and M.G.A. have no conflict of interest. R.P.P has served as a consultant and was previously employed by ViTToria Biotherapeutics. M.R. holds multiple patents related to CAR T-cell immunotherapy. M.R. has served as a consultant for Bristol Myers Squibb, GLG, Guidepoint, Lumicks, Acyla Therapeutics, Modex, and AbClon. M.R. receives research funding from AbClon, LUMICKS, Beckman Coulter, and Oxford Nano Imaging. M.R. is the scientific founder of viTToria Biotherapeutics which has licensed part of the technologies described in this paper.

### Funding

- Hematology Clinical Research Training Program, National Heart, Lung, and Blood Institut: T32 HL07439
- American Society of Hematology Research Training Award for Fellows
- Lymphoma Research Foundation
- American Society of Hematology Graduate Hematology Award
- National Cancer Institute: R37-CA-262362-02
- Leukemia and Lymphoma Society
- Gilead Research Scholar Award in Hematology
- Emerson Collective
- Laffey McHugh Foundation
- Parker Institute for Cancer Immunotherapy
- Berman and Maguire Funds for Lymphoma Research at Penn

