## Supplementary material for "Harnessing the CD2 axis to broaden and enhance the efficacy of CAR T cell therapies": Suppl figures

#### SUPPLEMENTAL METHODS

##### *Cell lines and primary samples*

Unless otherwise specified, all cell lines were cultured in R10 media (Roswell Park Memorial Institute medium 1640 (RPMI; Gibco; Cat#11875-085) supplemented with 10% fetal bovine serum (FBS, Gibco; Cat# 16140-071), 1% penicillin and 1% streptomycin (Gibco; Cat# 15140-163), 1% GlutaMAX supplement (Gibco; Cat# 35050-079, and 1% HEPES (Gibco; Cat# 15630-130)) in a 37°C incubator with 5% $CO_2$. All cell lines were authenticated by short tandem repeat (STR) analysis and tested for mycoplasma using a MycoAlert Plus Mycoplasma Detection Kit (Lonza; Cat# LT07-710). Nalm6, Jurkat, OCI-Ly18, and HEK293T cell lines were purchased from American Type Culture Collection (ATCC). TH20, a primary patient-derived xenograft model of T-cell acute lymphoblastic leukemia (T-ALL), was obtained as previously described (39). All cell lines were transduced with a lentivirus encoding click beetle green (CBG) and green fluorescent protein (GFP). Primary Sezary cells were provided by the clinical practices of Dr. Alain Rook. Primary acute myelogenous leukemia (AML) cells and other primary T-ALL cells were collected from patients treated at the Hospital of the University of Pennsylvania, as per IRB Protocol #855418.

##### *CD2 short guide RNA (sgRNA) optimization*

CRISPR sgRNAs were designed using *Benchling* (https://www.benchling.com). For *CD2*, *CD58,* and the *murine CD2*, sgRNA sequences were designed to target early exon sequences (**Table S1 and Table S2**), and *in vitro* transcribed using the GeneArt Precision gRNA Synthesis Kit (Invitrogen; Cat# A29377) for screening. Cells were electroporated using the Lonza 4D-Nucleofector Core Unit. Primary T-cells (human or mouse) were electroporated using the P3 Primary Cell 4D-Nucleofector X Kit L (Lonza; Cat# V4XP-3024). For Cas9 and sgRNA delivery, the ribonucleoprotein (RNP) complex was initially formed by incubating 10 µg of TrueCut Cas9 Protein v2 (Lonza; Cat# A36499) with 10 µg of sgRNA for 10 minutes at room temperature. $5x10^6$ cells were spun down at 300 × *g* for 5 minutes and resuspended in 100 µL in the specified buffer. The RNP complex and 100 µL of resuspended cells were combined and electroporated using pulse code EO-115. After electroporation, human T-cells were incubated in standard media containing 20 ng/mL of supplemental cytokines IL-7 and IL-15 at a concentration of $2×10^6$ cells/mL at 37°C. For mouse T-cells were used mouse-IL7 and human-IL2 at a concentration of 2×106 cells/mL at 37°C. CD2 or murine CD2 expression was subsequently monitored at each of the indicated days. After initial CD2 sgRNA screening, all subsequent experiments were performed using CD2gRNA8 for the human CD2 and CD2gRNA5 for mouse CD2.

##### *Lentivirus production*

Replication-defective, third-generation lentiviral vectors were produced using HEK293T cells. Approximately $8x10^6$ cells were plated in T150 culture vessels in standard culture media and incubated overnight at 37°C. 18-24 h later, cells were transfected using a combination of Lipofectamine 2000 (116 µL, Invitrogen; Cat#11668-019), pVSV/G or pCocal (7 µg), pRSV/Rev (18 µg), pGag/Pol (18 µg) packaging plasmids and 15 µg of expression plasmid. Lipofectamine and plasmid DNA were diluted in 4 mL Opti-MEM media (Gibco; Cat# 31985-070) before transfer into lentiviral production flasks. At both 24 and 48 h following transfection, culture media was isolated and concentrated using high-speed ultracentrifugation (8,500 rpm overnight or 25,000 rpm for 2.5 hours).

CARs targeting CD2 were generated based on the antibody variable region sequences of monoclonal antibodies MEDI507, OKT11, T11-2, and TS2 18.1.1. Single-chain variable fragments (scFv) were designed in both orientations (from variable light chain to variable heavy chain and vice versa) with three glycine-serine-serine-serine linkers and synthesized at GenScript Biotech (**Table S3**). After initial CD2 screening, all experiments were performed using MEDI507 in a light-to-heavy chain orientation. All CAR constructs were composed of an scFv, 4-1BB costimulatory domain, and CD3ζ costimulatory domain, unless otherwise noted.

##### *Manufacturing of primary human genome-engineered CAR T-cells*

Human T-cells were procured through the University of Pennsylvania Human Immunology Core (IRB #705906). $CD4^+$ and $CD8^+$ cells were combined at a 1:1 ratio and used for electroporation. CRISPR-Cas9 sgRNAs were generated using *in vitro* transcription, as described above. Mock KO cells were electroporated using the same procedure as described without the presence of an RNP complex. After electroporation, T-cells were incubated at 37°C for 24 hours and subsequently activated using CD3/CD28 Dynabeads (Gibco; Cat# 40203D) at a ratio of 3 beads/cell. The following day, CAR lentiviral vectors were added to stimulated cultures at a multiplicity of infection (MOI) of 1. Beads were removed at Day 5 of stimulation, and cells were counted every 2-3 days using a MoxiGO II (Orflo) until growth kinetics and cell size demonstrated they had rested from stimulation. All T-cells were initially grown with 20 ng/mL of supplemental cytokines IL-7 and IL-15 that were decreased to 0 ng/mL by the end of the expansion.

***General multiparameter flow cytometry***
Cells were resuspended in FACS staining buffer (phosphate buffered saline (PBS)+2% FBS) using one or more of the following antibodies: CD2 (UCHT2, 17H2L, or BL1a), CD4 (OKT4), CD8 (SK1), CD45RA (2H4, ALB11), CCR7 (G043H7), PD1 (EH12.2H7), LAG3 (7H2C65), CD3 (UCHT1, OKT3), hCD45 (J33), mCD45 (30-F11), PD-L1 (29E.2A3), mouse CD2 (RM2-5), mouse PD-1 (EH12.2H7), CAR2 and murine CAR19 were detected by incubating cells with an anti-G4S linker antibody conjugated to PE (CST; Cat# 71645S). Human CAR19 was detected using an anti-idiotype antibody provided by Novartis Pharmaceuticals. All changes in overall tumor or T-cell counts reflected by absolute cell counts were determined using Flow-Count Fluorospheres (Beckman; Cat# 754053). Cell viability was established using ViaKrome 808 Fixable Viability Dye (Beckman; Cat# C36628) or LIVE/DEAD Fixable Violet Dead Cell stain (Invitrogen; Cat# L34955), and data were acquired on a CytoFlex LX Flow Cytometer (Beckman). All data analysis was performed using FlowJo 10.8.0 software (FlowJo, LLC).

***Bioluminescence-based cell survival in vitro assays***
Cell lines (Jurkat, Nalm6, and OCI-Ly18) were engineered to express CBG, and cell survival was measured using bioluminescent quantification. D-luciferin potassium salt (Gold Biotechnology; Cat# 115144-35-9) was added to cell cultures (final concentration 15 μg/mL) and incubated at 37°C for 10 min. Bioluminescent signal was detected using a BioTek Synergy H4 imager, and signal was analyzed using BioTek Gen5 software. Percent cytotoxicity was calculated using a control of target cells without effectors.

***Flow cytometry-based cell survival in vitro assays***
Primary T-ALL, AML, or Sezary cells were stained with CellTrace Violet (CTV; Invitrogen; Cat# C34557) prior to plating with CAR T-cells or control untransduced T-cells. Reagents were used according to the manufacturer's protocol. After 48 h, cells were stained with ViaKrome 808 Fixable Viability Dye (Beckman; Cat# C36628) and analyzed by flow cytometry (CytoFlex LX; Beckman) to determine the absolute count of ViaKrome 808-negative and $CTV^+$ cells. Absolute cell counts were determined using Flow-Count Fluorospheres (Beckman; Cat# 754053). Percent cytotoxicity was calculated using a control of target cells without effectors.

***CellTrace Violet proliferation assays***
T cells were resuspended in PBS containing CellTrace Violet (CTV; 1:1,000 dilution) at a concentration of $1 \times 10^6$ cells per milliliter and incubated for 15 minutes at 37 °C. Cells were then washed with 10 mL of R10 medium, resuspended, and plated in co-culture with cancer cells at an effector-to-target (E:T) ratio of 0.25:1. After 5 days, proliferation was assessed by flow cytometry. The absolute number of $CD3^+$ T cells was determined using counting beads.

***Mitochondrial Respiration (Agilent Seahorse XF)***
Mitochondrial respiration was assessed using the Agilent Seahorse XF96 extracellular flux analyzer. XF96 wells were pre-coated with Cell-Tak (Corning) following the manufacturer's instructions, incubated overnight at 37°C, air-dried, and stored at 4°C. Prior to assay, XF RPMI medium (non-buffered) supplemented with 10 mM glucose, 2 mM L-glutamine, and 5 mM HEPES was prepared fresh and adjusted to pH 7.4. T cells were centrifuged at 500 × g for 5 minutes, washed with PBS, and resuspended in XF assay medium. A total of $1 \times 10^5$ cells per well were plated. The plate was centrifuged at 1,000 × g for 2 minutes and incubated at 37°C in a non-$CO_2$ incubator for 15 minutes. Baseline oxygen consumption rate (OCR) and extracellular acidification rate (ECAR) were measured, followed by sequential injections of 1.5 μM oligomycin A, 2.5 μM BAM15, and 0.5 μM rotenone with 0.5 μM antimycin A. ATP production rates were calculated using the Seahorse XF T Cell

Metabolic Profiling Report Generator, which distinguishes mitochondrial ATP production (mitoATP, pmol ATP/min) from glycolytic ATP production (glycoATP, pmol ATP/min).

***Cell-to-cell avidity experiments***

Z-Movi-compatible acoustofluidic chips were coated with poly-L-lysine for 3h prior to attaching a monolayer of tumor cells. CellTrace Far-Red (Invitrogen; Cat# C34564) labelled CAR T-cells were allowed to incubate on the monolayer of tumor cells for 15 min, and then a ramping acoustic force was applied. Cell-detachment was analyzed using ImageJ and R. Avidity experiments were conducted according to the manufacturer's (LUMICKS$^{TM}$) instructions and recommendations.

***Glass-supported planar lipid bilayer confocal imaging***

Supported lipid bilayers were prepared by fusing small liposome droplets on clean glass coverslips as previously described(41). Briefly, the liposome was trapped in a μ-Slide VI 0.4 chamber (Ibidi, Germany, Cat# 50-305-784). Lipid bilayers were first blocked with 5% casein for 20 min and then incubated with 6.3 nM of streptavidin (Thermo Fisher) for 15 min. After washing with imaging buffer (HEPES-buffered saline), lipids were incubated with biotinylated CD19 protein (R&D Systems, Cat# 9269-CD) conjugated with AlexaFluor 488, respectively, at room temperature for 30 min. After washing with the imaging buffer, lipid bilayers were blocked with 2.5 μM of D-biotin to saturate the streptavidin-binding sites. CAR T-cells were then stimulated on lipid bilayers for 30 min at 37°C. Cells were then fixed in 4% paraformaldehyde for 10 min and then permeabilized in 0.5% Triton-X100 and 10% normal donkey serum in PBS for 30 min at room temperature. Then, the cells were stained with primary antibodies against CD2 (ThermoFisher, Cat# 914-MSM3-P1ABX) and phosphorylated CD3 zeta (Abcam, Cat# ab68236) or CD19 (R&D Systems, Cat# AVI9269). An Olympus FV3000 confocal microscope with 60x (NA 1.35) oil objective was used to obtain confocal image data. The fluorescence intensity is calculated as the sum of fluorescence at the synapse for individual cells.

***Confocal imaging immune synapse with Leica Stellaris***

Prior to the experiment, 18-well Ibidi chambers (81817, Ibidi) were covered with Poly-L lysine (P4707, Sigma) for 15mins at RT and washed 3X in PBS. Afterwards, 200 000 NALM6-GFP cells per well were seeded in RPMI supplemented with 2% FBS for 30 minutes to promote cell attachment to the chamber. Tumour cells were then co-cultured for 20 minutes with CD2 WT and KO CART19 at a 1:1 ratio of CAR-positive to target cells. After the co-cultures, the samples were washed with PBS and fixed with 4% PFA (J19943.K2, ThermoFisher) for 15 minutes at room temperature. After fixation, the cells were washed 3x with PBS, blocked with PBS + 5% BSA (A7906, Sigma) for 30 mins at RT, and stained with mouse anti-human CD5 (UMAB9, ThermoFisher) at 1/100 dilution in blocking buffer for 1 hr at RT. After surface staining, samples were washed 3x with PBS and permeabilized with cold methanol for 10 mins at RT and blocked with PBS + 5% BSA for 30mins at RT. After blocking, samples were stained with rabbit anti-human pCD3z (ab68235, Abcam) overnight at 4°C. Then, the samples were washed 3x with PBS and stained with goat anti-mouse AF568 (A1104, ThermoFisher) and goat anti-rabbit AF647 (A21245, ThermoFisher) both at 1/500 dilution in blocking buffer for 1 hr at RT. The samples were washed 3x with PBS and imaged on a Leica Stellaris with a 63x/1.4 Oil objective.

***In vivo immunodeficient mouse models***

6-10-week-old NOD-SCID-$\gamma c^{-/-}$ (*NSG*) mice were obtained from the Jackson Laboratory and maintained in pathogen-free conditions. All target cells were engineered to express CBG. Animals were injected via tail vein with $1x10^6$ Jurkat, TH20, Nalm6, or Nalm6-PDL1-overexpressing cancer cells in 0.15 mL sterile PBS. For scRNAseq experiments, 5.0 x $10^6$ OCI-Ly18 were subcutaneously injected. After engraftment, $1x10^6$ T-cells ($CAR^+$ or an equal number of UTD control) were injected via tail vein in 0.15 mL sterile PBS, unless otherwise indicated. Animals were monitored for signs of disease progression and overt toxicity, such as xenogeneic graft-versus-host disease, as evidenced by >20% loss in body weight, loss of fur, diarrhea, conjunctivitis, and disease-related hind limb paralysis. Disease burdens were monitored over time using a Xenogen IVIS bioluminescent imaging system or the Lumina S5 imaging system for tumor flux.

***Single cell RNA sequencing and analysis***

Peripheral blood was collected from all mice, followed by red blood cell lysis. Human cells were enriched using the EasySep Mouse/Human Chimera Isolation Kit (StemCell Technologies; cat. no. 19849) according to the manufacturer's instructions. To further purify T cells, enriched samples were stained with anti-human CD3, CD5, and CD2 antibodies (APC, Biolegend) and sorted using a BD FACSMelody Cell Sorter. Prior to single-cell capture, cells were barcoded using TotalSeq-A Hashtagging Antibodies (BioLegend) to allow for sample multiplexing. After sorting, cells were washed twice with 0.04% BSA in PBS and loaded onto the 10X Chromium Controller. Libraries were prepared using the 10X Genomics v2 Chemistry kit, according to the manufacturer's instructions. Next-generation sequencing on libraries was performed using the Illumina NovaSeq 6000. Raw single-cell RNA-seq data were processed using Cell Ranger software (v5.0.1; 10x Genomics), with transcript alignment and mapping performed against the GRCh38 human reference genome. Filtered gene expression data were processed in Seurat v 5.3.0 (42). Quality control filters were applied to retain cells with >200 and <5,500 detected RNA features and <10% mitochondrial gene content. Data were normalized and scaled using standard Seurat workflows. Dimensionality reduction was performed with principal component analysis by using a curated list of canonical T cell genes described by Szabo et al. to drive clustering (43). The top 30 principal components were used to drive unsupervised clustering via UMAP at a resolution of 0.6. Differential gene expression between clusters across experimental groups was assessed using the FindMarkers function, which applies a two-sided Wilcoxon rank-sum test followed by Benjamini–Hochberg correction for multiple comparisons. For pathway analysis, we used clusterProfiler v4.0 (44).

***CD2 patient validation cohorts***

The flow cytometry archives (2002-2014) at the Children's Hospital of Philadelphia (CHOP) were reviewed to identify cases of T-ALL in the pediatric and young adult age group (0-25 years). Diagnosis of T-ALL was confirmed per WHO criteria(45). Archived flow cytometric histograms that were saved as PDFs were reviewed. Flow cytometric data were collected using Beckman Coulter FC 500 Series flow cytometers in a College of American Pathologists (CAP) accredited clinical flow cytometry laboratory. Bead controls, titration of newer antibody lots, and daily comparison of instruments were performed. One million cells/microliter single cell suspensions of bone marrow, peripheral blood or tissue were incubated with 5 color antibody panels including CD45-PE-Cy7, CD4-PE, CD8-FITC, CD7-PE-Cy5.5, CD3-ECD, CD2-FITC, CD56-PE, B-cell, myeloid markers, and isotype control antibodies (Beckman Coulter, CA). Twenty thousand events were collected. Data analysis was performed using FCS Express (De Novo software, CA). Negative gates were based on isotype controls. Immature blasts were identified using CD45 vs. SSC gating. T-cell markers CD2, sCD3, CD5, and CD7 were assessed on CD45dim blasts. Normal CD45bright mature T-cells served as controls for assessment of antigen.

The phenotypic profile of peripheral T-cell lymphomas from 91 patients diagnosed and treated at the European Institute of Oncology was analyzed via immunohistochemistry for CD2, CD3, CD5, and CD7 expression by a board-certified hematopathologist. CITE-seq data from patients enrolled in AALL0434 were obtained from Xu J, *et al.*(46). AALL0434 (NCT00408005) was a Children's Oncology Group (COG) phase 3 clinical trial that enrolled subjects from January 22, 2007 until July 25, 2014; trial results were previously published(47). T-ALL subjects enrolled on a companion biology study used for sample banking, AALL03B1 (NCT00482352) from January 22, 2007 until August 08, 2010 or AALL08B1 (NCT01142427) from August 9, 2010 until July 25, 2014. AALL03B1, AALL08B1, and AALL0434 were approved by the NCI Cancer Evaluation and Therapeutic Program (CTEP), local IRBs at participating centers, and the Pediatric Central Institutional Review Board (IRB). Informed consent was obtained from all study participants and/or their legally authorized representative in accordance with the Declaration of Helsinki. Genomic studies were approved by the local IRB at the Children's Hospital of Philadelphia, COG, and CTEP. Samples were decoded and assigned a unique study identifier (USI). Samples were banked at the COG biorepository at Nationwide Children's Hospital in Chicago, IL. CITE-seq data were normalized using CLR normalization across cells (margin = 2). Original cell annotations, based on a 7-step process involving a projection to healthy pediatric thymus and bone marrow developmental reference, were retained. Data was integrated using the *IntegrateLayers* function in Seurat v5 using reciprocal PCA and visualized on a UMAP using the top 30 principal components(48). Surface and RNA expression profiles were plotted using the *FeaturePlot* and *VinPlot* functions in Seurat v5 with default parameters. All code generated for processing of the CITE-Seq are available at: https://github.com/tanlabcode/SC_TALL. Surface and RNA expression for CD2 and other T-cell markers were plotted in n=16,199 healthy thymocytes from 3 pediatric donors and visualized using the *VinPlot* function in Seurat v5.

***CD58 immunohistochemistry of primary B-cell lymphomas***

CD58 immunohistochemical stain was performed on selected patient samples. Five-micron sections of formalin-fixed paraffin-embedded tissue were stained using antibody against CD58 at 1:100 dilution (R&D systems, Cat# AF1689). Staining was done on a Leica Bond-IIITM instrument using the Bond Polymer Refine Detection System (Leica Microsystems DS9800) and Goat IgG VisUCyte HRP Polymer (R&D Systems, Cat# VC004-25). Heat-induced epitope retrieval was done for 20 minutes with ER2 solution (Leica Microsystems AR9640). All the experiments were performed at room temperature. Slides were washed three times between each step with bond wash buffer or water. Positive staining was visualized and scoring of percentage of tumor cells and intensity of staining were determined by at least two hematopathologists. Hematopathologists were blinded to patient outcomes and categorization at the time of immunohistochemical evaluation. The percentage of tumor staining was determined by evaluating previous H&E and immunohistochemical stains performed at initial diagnosis. Positive staining was defined as clear membranous staining present. Intensity was determined by evaluating immunohistochemical stain controls, with the strongest intensity membranous staining interpreted as 3+ and the weakest intensity staining observed interpreted as 1+ staining. Variable or non-specific staining was scored as negative. The final H-score was calculated by multiplying the intensity of staining by the percentage of tumor cells staining.

##### *Western blot and phospho-LCK analysis*

Protein extracts were obtained from T cells following 20 minutes of stimulation on plates coated with recombinant human CD19 (catalog #CD9-H52H2), PD-L1 (catalog #PD1-H52H3), or CD58 (catalog #LF3-H5225 proteins. 1500 ng of each protein was coated/well. Cells included CD2 wild-type or CD2 knockout CART19, as well as CD2 knockout CART19 expressing either the full PD-1:CD2 switch receptor or its truncated version lacking the intracellular CD2 domain. Cells were lysed in RIPA buffer (Research Products International, #R26200-250.0) supplemented with protease and phosphatase inhibitor cocktails (Cell Signaling Technology, #5871S and #5870S). Protein concentrations were determined using the BCA assay, and 20 µg of total protein per sample were resolved by SDS-PAGE on 4–20% Mini-PROTEAN TGX Precast Protein Gels (Bio-Rad, #4561094), then transferred onto 0.2 µm nitrocellulose membranes. Membranes were blocked using Intercept (PBS) Blocking Buffer (LI-COR Biosciences, #927-70001) and incubated overnight at 4 °C with primary antibodies against phospho-LCK (Tyr505, Cell Signaling Technology, #2751S) and β-actin (Cell Signaling Technology, #3700S) as a loading control. After washing, membranes were incubated at room temperature with appropriate IRDye-conjugated secondary antibodies (LI-COR Biosciences, Goat anti-Rabbit IgG, #926-32211; Goat anti-Mouse IgG, #926-68052), and scanned using an Odyssey CLx imaging system (LI-COR). Band intensities were quantified using ImageJ software, and phospho-LCK levels were normalized to β-actin for each sample.

##### Jurkat triple-reporter gene assay

Jurkat expressing three inducible reporter genes: NFAT-GFP, NF-κB-CFP and AP-1-mCherry were transduced with our CAR constructs +/- switch full-length or truncated receptor. Target cells were stained with CellTrace Violet (#C34557; Thermo Fisher Scientific), while Jurkat reporter cell lines were labeled with CellTracker Deep Red (#C34565; Thermo Fisher Scientific) following manufacturer's protocols. Co-cultures of target cells and CAR triple reporter Jurkat cell lines were incubated at a 1:1 effector-to-target ratio with 50,000 cells per cell type. After 24 hours at 37°C, co-cultures were harvested and analyzed for GFP, CFP and mCherry reporter gene expression, measuring the geometric median fluorescence intensity ratio of each fluorescent reporter.

##### *Murine CRISPR–Cas9-edited CAR T-cell manufacturing and in vivo treatment*

Dynabeads (Thermo Fisher) at a 2:1 bead-to-cell ratio in R10 medium supplemented with 0.05 mM 2-mercaptoethanol, 10 ng/mL murine IL-2, and 10 ng/mL murine IL-7. After 24 hours of activation, CD2 was knocked out using CRISPR–Cas9 ribonucleoprotein (RNP) electroporation with a CD2-targeting gRNA (CD2gRNA5, sequence provided in **Table S2**), using the Lonza 4D Nucleofector and pulse code DN-100. Control cells were mock-electroporated without sgRNA. Immediately following electroporation, cells were incubated for 1 hour at 37 °C, and then transferred into RetroNectin-coated plates pre-loaded with a retroviral vector encoding a murine anti-CD19 CAR (muCART19) containing a 4-1BB costimulatory domain and CD3ζ signaling domain. For groups expressing the PD-1:CD2 switch receptor, a bicistronic retroviral vector encoding muCART19 linked via a T2A sequence to a murine PD-1:CD2 switch receptor. Retroviral transduction was performed by spinfection (1,000 g, 1 h, 32 °C). Cells were cultured for 5–6 days post-transduction with fresh cytokines every 48 hours. On day 5 or 6, Dynabeads were removed magnetically. Transduction efficiency ($CAR^+$ and $PD\text{-}1^+$ for conditions expressing the switch receptor) and CD2 knockout efficiency were assessed by flow cytometry. Cells were then washed and resuspended in PBS for intravenous injection. For in vivo experiments, 6–8-week-old female BALB/c mice were injected intravenously with 1.0 × $10^6$ A20 lymphoma cells. On day 3 post-tumor injection, mice received cyclophosphamide (CTX, 200 mg/kg intraperitoneally) for lymphodepletion. On day 4, mice were treated with 1.0 × $10^6$ total murine T cells, corresponding to one of the following groups: untransduced T cells (UTD), $CD2^{WT}$ muCART19, $CD2^{KO}$ muCART19, or $CD2^{KO}$ muCART19 expressing the PD-1:CD2 switch receptor. Tumor growth was monitored by caliper measurements every 2–3 days, and volume was calculated using the formula (length × $width^2$) / 2. Mice were euthanized upon reaching humane endpoints, and survival was recorded (41).

##### *Syngeneic mouse model*

Six- to eight-week-old female BALB/c mice were purchased from The Jackson Laboratory and housed under conventional conditions (BSL-1). To establish the syngeneic A20 lymphoma model, 2 × $10^6$ A20 cells were resuspended in 100 µL of a 1:1 mixture of PBS and Matrigel (Corning) and injected subcutaneously into the right flank. Fourteen days after tumor implantation, mice were treated with 200 mg/kg of cyclophosphamide via intraperitoneal injection for lymphodepletion. On day 15 post-implantation (when tumors reached approximately 100–200 $mm^3$), mice were randomized and treated with 1 × $10^6$ T cells via intravenous injection. Treatment groups included untransduced T cells (UTD), $CD2^{WT}$ murine CAR19 T cells, $CD2^{KO}$ murine CAR19 T cells, and $CD2^{KO}$ murine CAR19 T cells expressing the murine PD-1:CD2 switch receptor. Tumor volume was monitored every 2–3 days by caliper measurements and calculated using the formula (length × $width^2$)/2. Mice were euthanized upon reaching humane endpoints in accordance with protocols approved by the University of Pennsylvania Institutional Animal Care and Use Committee (IACUC).

##### *Statistical analysis*

Data were visualized and analyzed using Prism 10 software (GraphPad). All results are represented as either individual values or as mean values ± standard error of the mean (SEM), unless otherwise noted. All comparisons between two groups were performed using two-tailed unpaired Student's t-test. Comparisons between more than two groups were performed by one-way analysis of variance (ANOVA) with Tukey correction for multiple comparisons. In analyses where multiple groups were compared at multiple time points/ratios, two-way ANOVA was performed. Survival data were analyzed using the Log-Rank (Mantel-Cox) test unless otherwise noted. The p values were denoted with asterisks as follows: * $p < 0.05$, ** $p < 0.01$, *** $p < 0.001$, **** $p < 0.0001$.

### Supplemental Figure 1

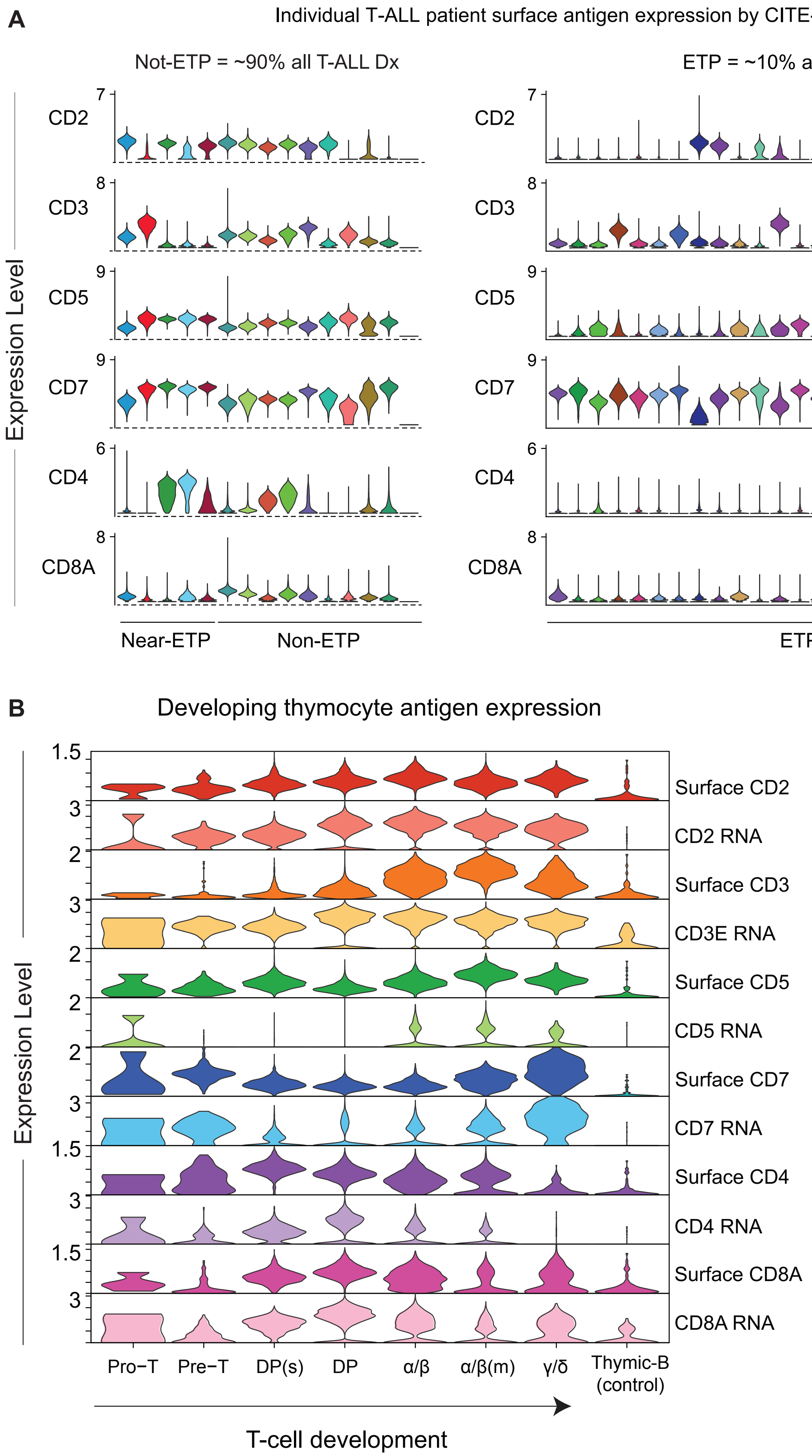

### Supplemental Figure 2

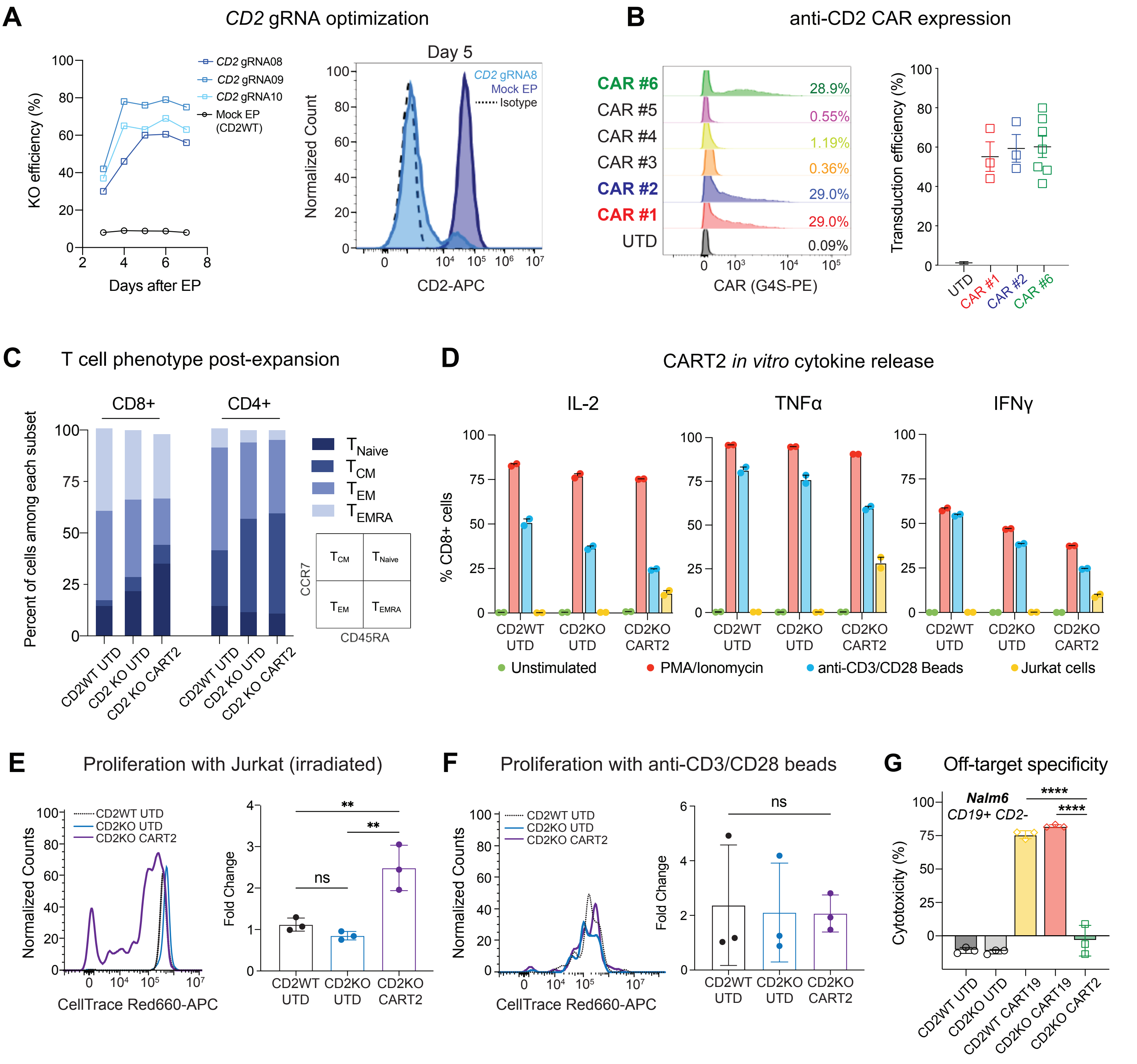

### Supplemental Figure 3

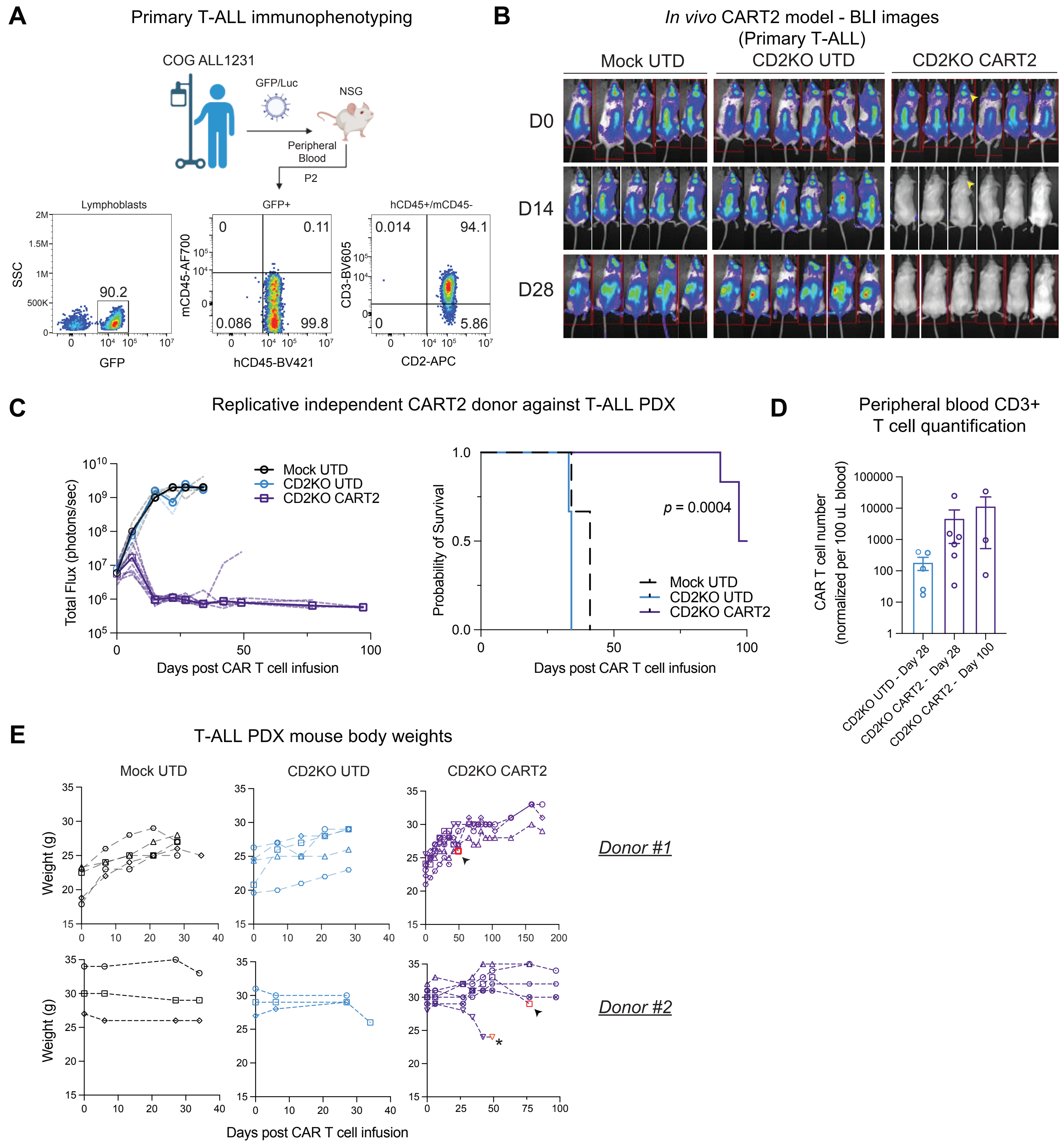

### Supplemental Figure 4

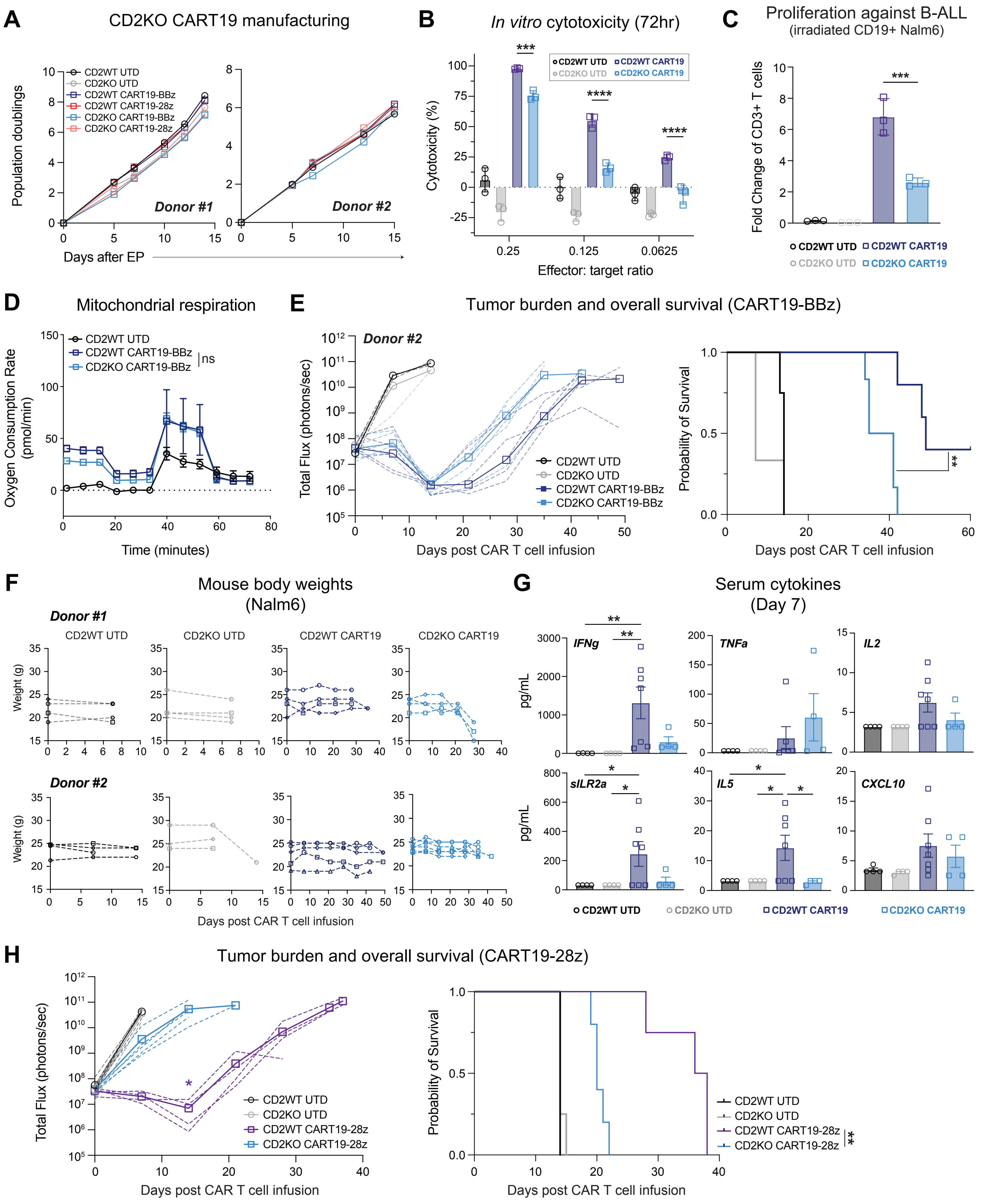

### Supplemental Figure 5

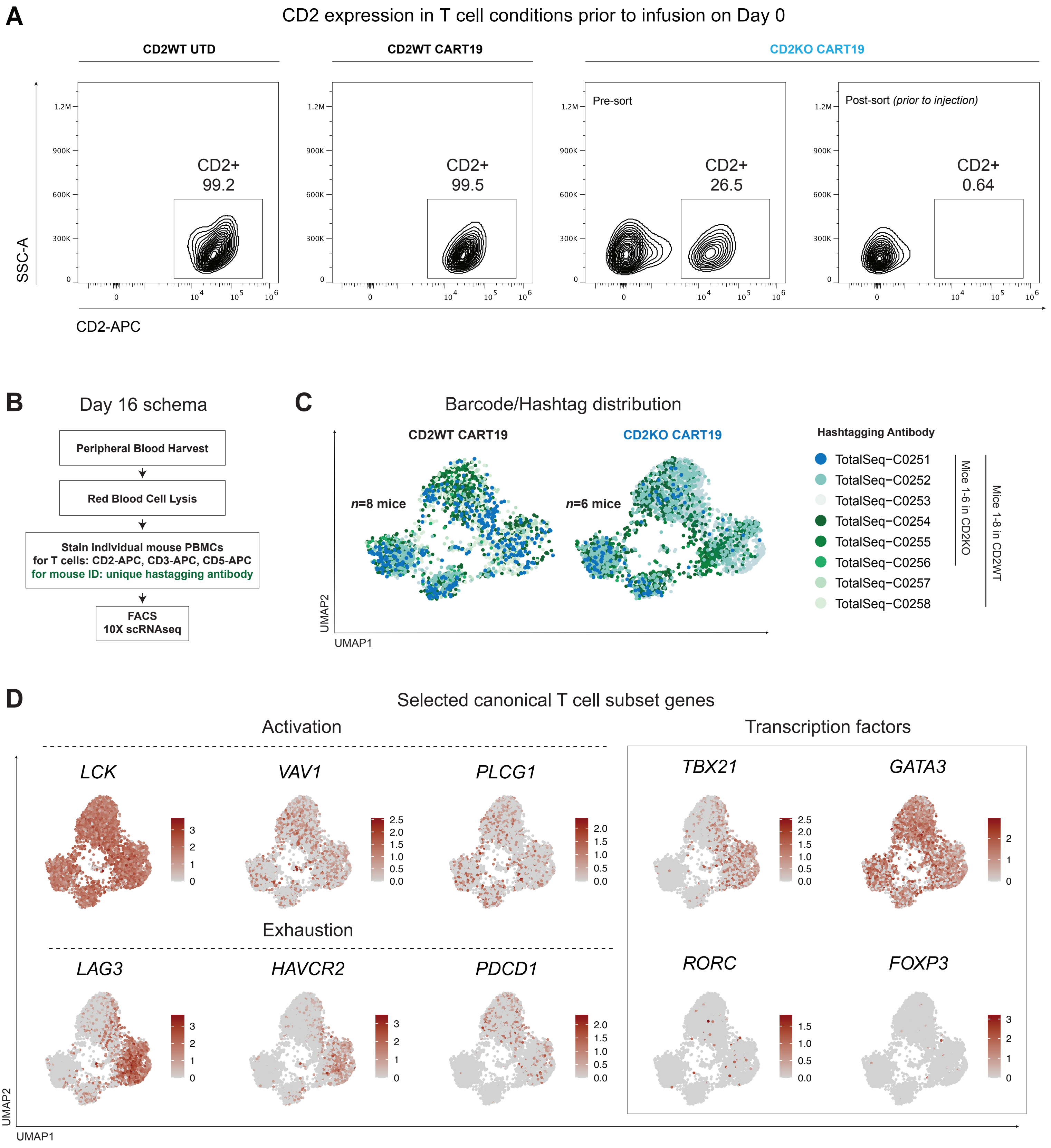

### Supplemental Figure 6

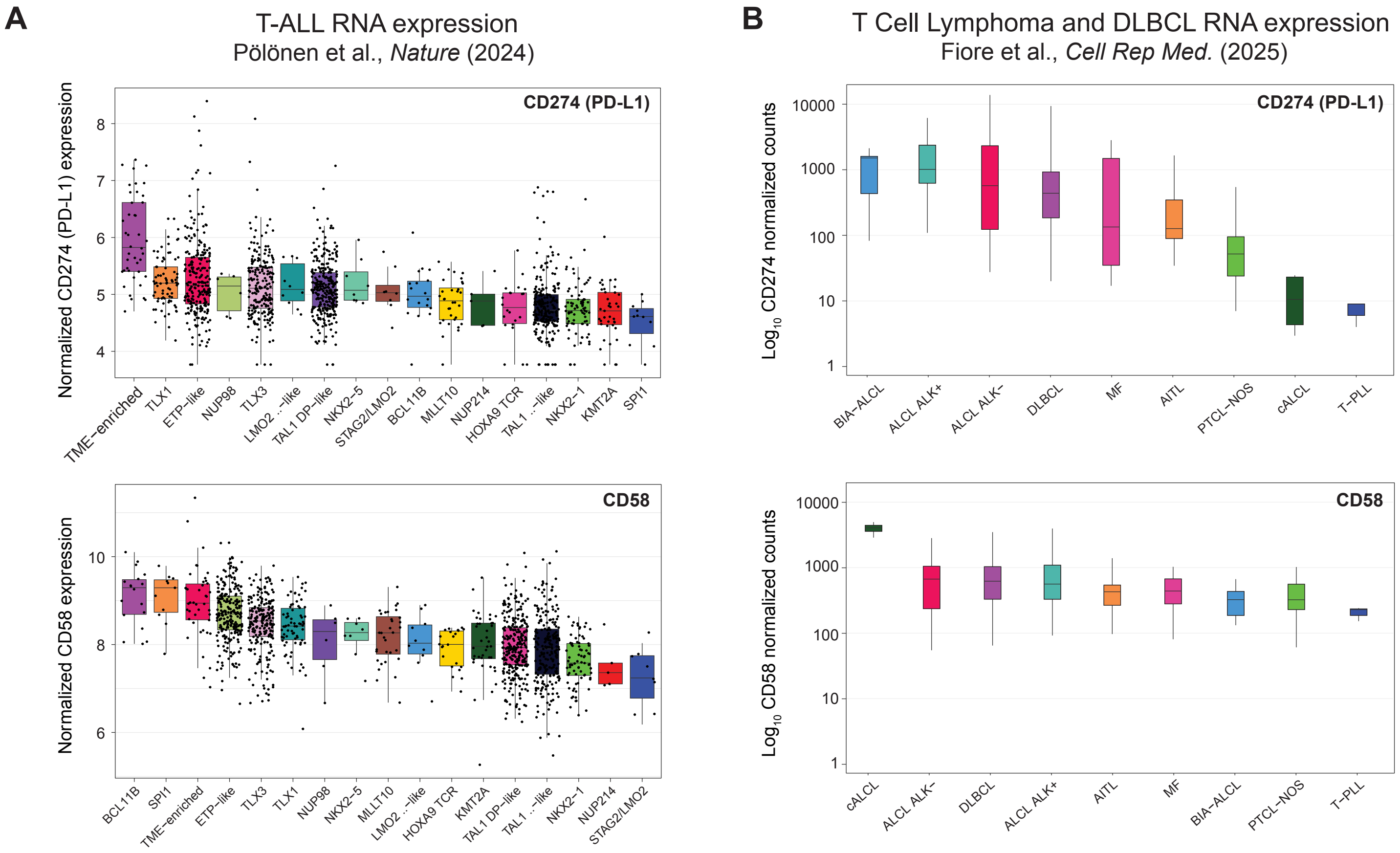

### Supplemental Figure 7

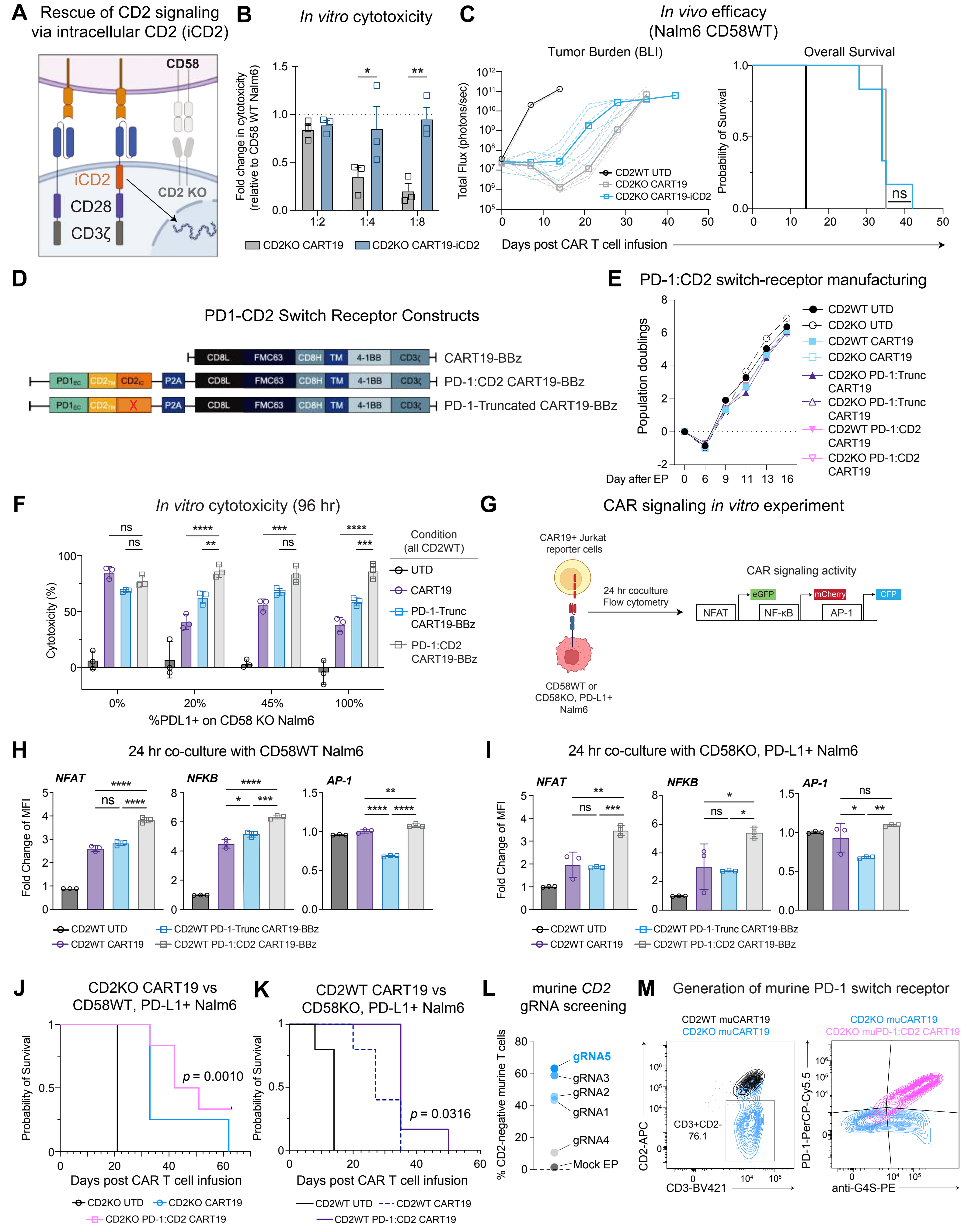

#### Full-length Western blots, associated with Fig 7D

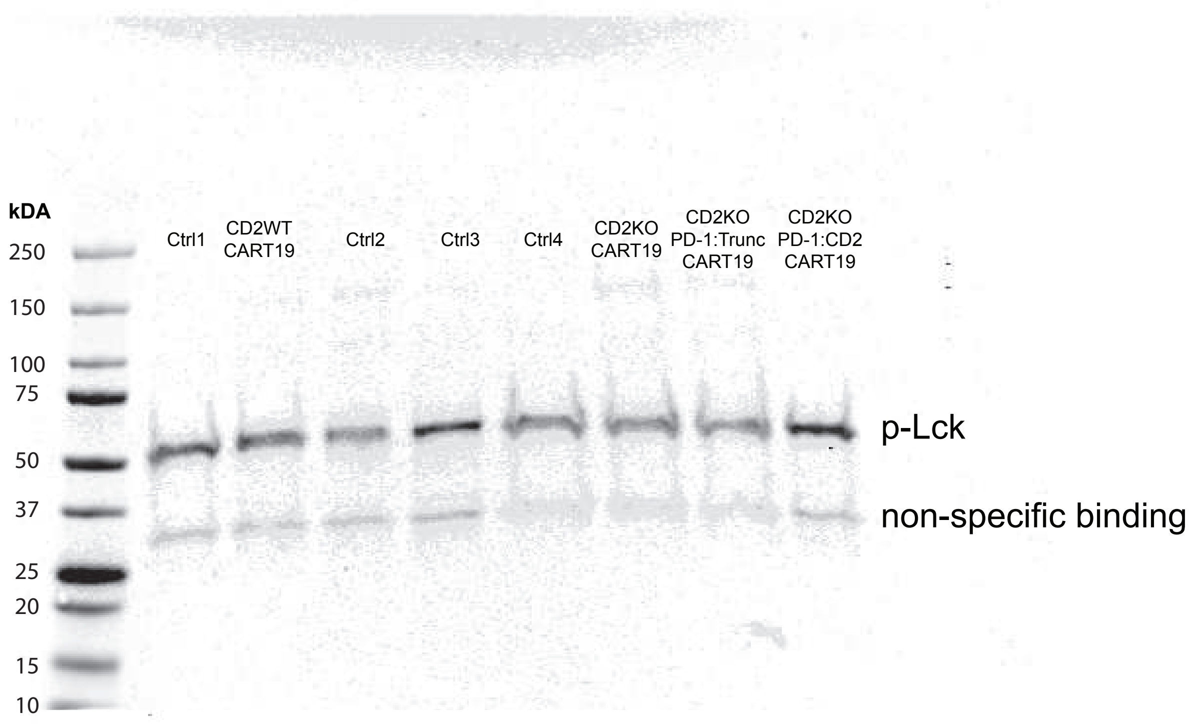


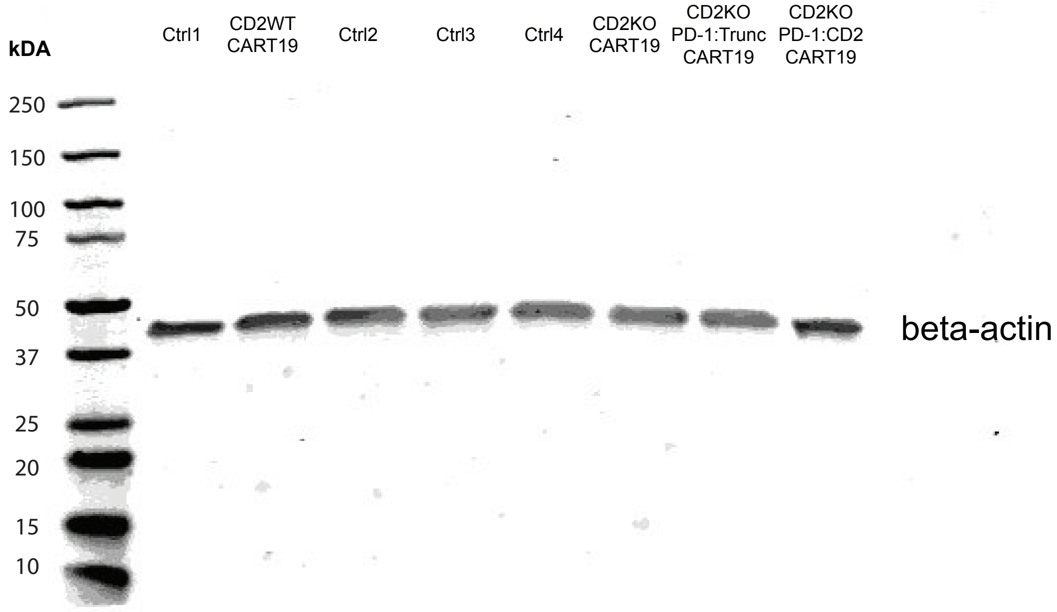
